# ETTIN-mediated auxin signalling is an angiosperm-specific neofunctionalization for carpel development

**DOI:** 10.1101/2024.09.02.610784

**Authors:** Aaron Chun Hou Ang, Sumanth Mutte, Dolf Weijers, Lars Østergaard

**Affiliations:** Crop Genetics Department, John Innes Centre, Norwich NR4 7UH, UK; Laboratory of Biochemistry, Wageningen University, Stippeneng 4, Wageningen, the Netherlands; Department of Biology, University of Oxford, South Parks Road, Oxford OX1 3RB, UK

**Keywords:** Evolution, angiosperms, non-canonical auxin signalling, neofunctionalization, gynoecium development

## Abstract

The phytohormone auxin affects processes throughout plant growth and development. While auxin signalling has been mainly attributed to a repressor degradation-based pathway, numerous alternative mechanisms for how auxin mediates its effect have been revealed in recent years. One such mechanism involves a direct auxin-induced switch in the transcriptional regulatory activity of the Auxin Response Factor (ARF) ETTIN (ETT). ETT lacks a conserved C-terminus domain involved in canonical pathway interactions but contains a middle region domain mediating auxin binding. As the ETT clade only exists in the angiosperms, it remains unknown when the pathway evolved. Here we provide evidence for a two-step origin of the ETT clade and its neofunctionalisation through the gain of auxin perception in gynoecium patterning. Phylogenetic analyses reveal the structural divergence of ETT and its paralogue ARF4 after their duplication from an ancestral euphyllophyte ARF3/4 clade. Auxin sensitivity was identified as an ETT-specific innovation that likely originated in the last common angiosperm ancestor. Furthermore, *in planta* complementation experiments demonstrated the full genetic redundancy of ETT and ARF4 in leaf and ovary development, but a specialised role for the ETT-mediated auxin signalling pathway in style development. Our work thus provides evidence that ETT was recruited from an ancestral role in leaf development and subsequently underwent neofunctionalisation through the acquisition of direct auxin sensing for a novel role in gynoecium patterning.

## INTRODUCTION

The transition of plants from their aquatic origin to a terrestrial habitat approximately 450M years ago is marked by numerous morphological innovations [1, 2]. Similarly, a key morphological innovation underlying the evolution of flowering plants (angiosperms) >120M years ago, is marked by the carpel and other floral organs that define the flowering plants (angiosperms). These novel organs were critical for the success and rapid radiation of this phylum [3, 4]. In angiosperms, numerous ancient gene families have greatly expanded, likely contributing to the evolution of their morphological complexity through the neofunctionalisation of duplicated paralogues [5–8]. This trend is also observed for the components of many phytohormonal signalling pathways, including the Auxin Response Factor (ARF) family of transcriptional regulators that mediate auxin signalling in plants [9, 10].

Auxin is well-known for its ubiquitous roles throughout plant development [11–13]. The ‘canonical’ pathway of auxin signalling posits that auxin-responsive gene expression is activated by the ARFs. In the absence of auxin, ARFs heterodimerise with the AUX/IAA transcriptional repressors through their C-terminal Phox/Bem1 (PB1) domains [14, 15], resulting in the recruitment of the TOPLESS (TPL) corepressor complex and gene repression [16–18]. Auxin acts as a molecular glue between its TIR1/AFB receptors and the AUX/IAA transcriptional repressors, ultimately leading AUX/IAA degradation and thus de-repression of ARF activity [19–21]. In addition to the canonical auxin signalling pathway, several alternative auxin signalling mechanisms have been described in recent years [22–26]. Furthermore, ARFs are divided into three major clades (A-, B- and C-ARF) that behave as either transcriptional activators or repressors [9, 27, 28]. In a simple bryophyte auxin signalling pathway, its A-ARF is the sole auxin-dependent activator while its B-ARF act as an auxin-independent competitor for target loci [29].

In the model angiosperm *Arabidopsis thaliana*, an atypical B-ARF, ETTIN (ETT – also known as ARF3) has been shown to mediate a novel auxin signalling pathway [25]. ETT lacks the PB1 domain necessary for canonical pathway interactions, but instead possesses a specific middle region (MR) that interacts directly with auxin and numerous protein families, including the TPL/TPR corepressors [30, 31]. Auxin binding to ETT disrupts the interaction between ETT and TPL, shifting ETT function to that of a transcriptional activator for the de-repression of ETT target genes. Many of these genes are important for gynoecium patterning and the loss-of-function of ETT leads to various gynoecium defects including reduced valve growth, medial and stigmatic tissue overgrowth, and a loss of radial style symmetry [32–35].

While the TIR1/AFB-dependent pathway is conserved in all land plants [9, 29], it is unknown when ETT-mediated auxin signalling evolved, considering the relatively recent origin of the ETT clade. ETT and its close paralogue, ARF4, emerged from a gene duplication event at the base of the angiosperms from an ancestral ARF3/4 clade [10, 36]. ETT and ARF4 are fully redundant in leaf dorsiventral polarity and mediolateral growth, but only partially redundant in gynoecium development [37, 38]. Considering the structural and functional differences between ETT and ARF4 in leaf and gynoecium development, it is possible that ETT-mediated auxin signalling arose as an angiosperm-specific innovation for carpel patterning while ETT and ARF4 retain the putative ancestral leaf development function of the ARF3/4 clade after their divergence.

Here, we show that the ARF3/4 clade emerged in an ancestral euphyllophyte and that ETT and ARF4 have undergone complex structural evolution in core angiosperm lineages. The ability of ETT to sense auxin emerged prior to core angiosperm radiation and is strongly linked to a portion of its MR domain. We demonstrate that the ARF3/4 clade acquired functions in leaf development in the seed plants while ETT and ARF4 acquired gynoecium-specific roles in the angiosperms. Notably, the auxin-sensitivity of ETT correlates with its ability to regulate style morphogenesis. Taken together, our data suggest that auxin sensing evolved as an ETT-specific neofunctionalisation from an ancestral leaf development function to mediate polarity establishment of the gynoecium of angiosperms.

## RESULTS

### The ARF3/4 clade emerged in the last euphyllophyte common ancestor

Previous studies on the evolutionary history of B-ARFs have yielded conflicting data regarding the origin of the ARF3/4 clade and its presence in the monilophytes [9, 10, 39, 40]. The discrepancies between studies might have arisen from the use of transcriptomic data for phylogenetically important clades that lacked genomic resources. To address this limitation, we utilised the recent availability of high-quality genomes of previously underrepresented clades (Table S1) for phylogeny construction of the B-ARFs (Figure 1A and S1).

**Figure 1.**
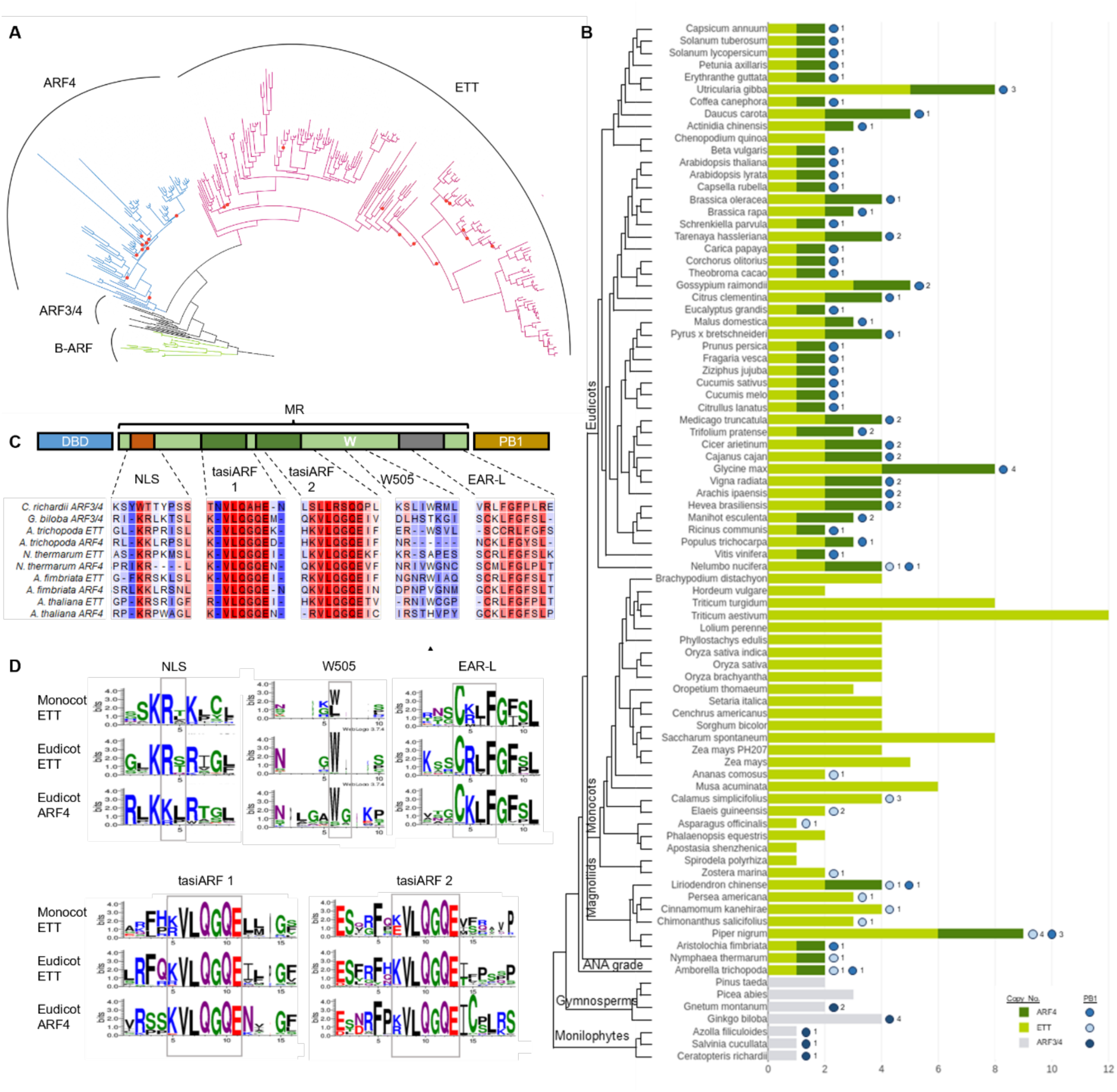
Phylogenetic analysis of the ARF3/4, ETT and ARF4 clades in the land plants. (A) Maximum likelihood phylogeny of the ARF3/4 clade. Red circles on the branches indicate poor bootstrap support (<75) for that split. (B) Gene copy number (bars) and PB1 domain absence/presence patterns (circles) of euphyllophyte ARF3/4, ETT and ARF4 orthologues. (C) Amino acid alignment of the middle region motifs in ARF3/4, ETT and ARF4 orthologues from species representing important euphyllophyte lineages. The black arrow highlights the position of the W505 residue. (D) Amino acid logos representing clade-enriched variants of conserved middle region motifs in eudicot and monocot ETT and ARF4 sequences.

In accordance with previous studies [9, 10], bryophyte and lycophyte lineages possess only a single B-ARF clade while extant gymnosperms share three distinct clades (ARF1/rest, ARF2, and ARF3/4) with the angiosperms (Figure S1). However, our phylogeny clusters a subset of Class B ARF sequences from the monilophytes *Ceratopteris richardii, Azolla filiculoides* and *Salvinia cucullata* with the ARF3/4 clade of the seed plants (Figure 1A and 1B), indicating that the ARF3/4 clade is present in monilophytes.

In angiosperms, two tasiARF binding sites are present in *ETT* and *ARF4* and binding by the *TAS3* tasiRNA to these sites inhibits *ETT* and *ARF4* expression [39] (Figure 1C and 1D). Mutations in these sites confer pleiotropic leaf and flower development phenotypes [41–43]. It was shown that the monilophyte *Ceratopteris pteridoides* ARF3/4 orthologue contained a single tasiARF site with a sequence that differs from those of seed plant orthologues [40]. In agreement with this finding, all monilophyte ARF3/4 orthologues included in our phylogeny lacked the canonical seed plant tasiARF site (Figure 1C and S2). These results suggest that the ARF3/4 clade originated in the last common ancestor of euphyllophytes but diverged functionally in monilophytes and seed plants.

### ETT and ARF4 have undergone divergent structural evolution in the angiosperms

The ARF3/4 clade diverged into the ETT and ARF4 clades in the last common ancestor of the angiosperms although ARF4 sequences were not found in Poaceae monocots [9, 10, 36]. No ARF4 orthologues could be identified from any of the monocot or Laurales genomes included in our phylogeny (Figure 1B). Conversely, we observe an increase in ETT copy number in many monocot genomes relative to that of the eudicots (Figure 1B). This suggests that ARF4 has been independently lost twice during angiosperm radiation while the ETT clade has expanded in the monocots.

A defining feature of ETT in the core eudicots is the lack of the PB1 domain that is important for ARF regulation by the TIR1/AFB-dependent auxin signalling pathway [10, 25, 31]. ETT orthologues of the ANA grade angiosperms and some magnoliids possess the full PB1 domain, but not in Poaceae monocots [10]. Through our alignment, we identify the presence of the full PB1 domain in ETT orthologues from species within the Alismatales, Asparagales, Arecales and Poales (Figure 1B), suggesting that the PB1 domain has been lost at least four times within the monocots rather than once in the common ancestor of all monocots.

As the PB1 domain is not a reliable indicator of ETT or ARF4 orthologue identity outside the core eudicots, we attempted to identify middle region motifs that might differ between the two clades through a machine learning approach (Figure 1C and 1D). While the NLS, tasiARF, EAR-L motifs and W505 residue were generally present in both ETT and ARF4 clades, clade-enriched variants were detected. The consensus ETT NLS sequence is KRx(K/R) while the ARF4 NLS sequence is KKxR. The EAR-L motif in eudicot ETT sequences is (C)RLFG while eudicot ARF4 sequences have the (C)KLFG variant, although both variants are found in monocot ETT sequences. The W505 residue is sometimes absent in monocot ETT sequences and is flanked by a LGA motif in eudicot ARF4 sequences. Within the ANA grade angiosperms, the W505 residue is present in the *Amborella trichopoda* ETT orthologue but missing in the ETT orthologue from *Nymphaea thermarum* (Figure 1C). These results indicate that while the general middle region structure is shared between ETT and ARF4 clades, the consensus sequence for important motifs have diverged between the two clades and motifs may be lost in different paralogues or orthologues.

### The *Amborella trichopoda* ETT orthologue exhibits direct auxin sensing

The *A. thaliana* ETT has been shown to bind auxin and the TPL/TPR corepressor proteins directly through its specialised middle region to mediate a novel auxin signalling pathway [25, 30, 31, 34]. While the ETT and ARF4 middle region motifs are generally well conserved (Figure 1C and 1D), it is unknown whether the ETT-mediated auxin signalling pathway is also present in other angiosperms. To assess the origin and conservation of ETT-mediated auxin sensing, we conducted a yeast-2-hybrid screen of the ETT, ARF4 or ARF3/4 orthologues from a monilophyte (*Ceratopteris richardii,* CriARF3/4), a gymnosperm (*Ginkgo biloba,* GbiARF3/4), two ANA grade angiosperms (*A. trichopoda*, AtrETT and AtrARF4; *N. thermarum*, NthETT and NthARF4), and a eudicot (*A. thaliana*, AthETT and AthARF4) with their respective TPL/TPR partners and tested the auxin sensitivity of interacting pairs (Figure 2 and S3).

**Figure 2.**
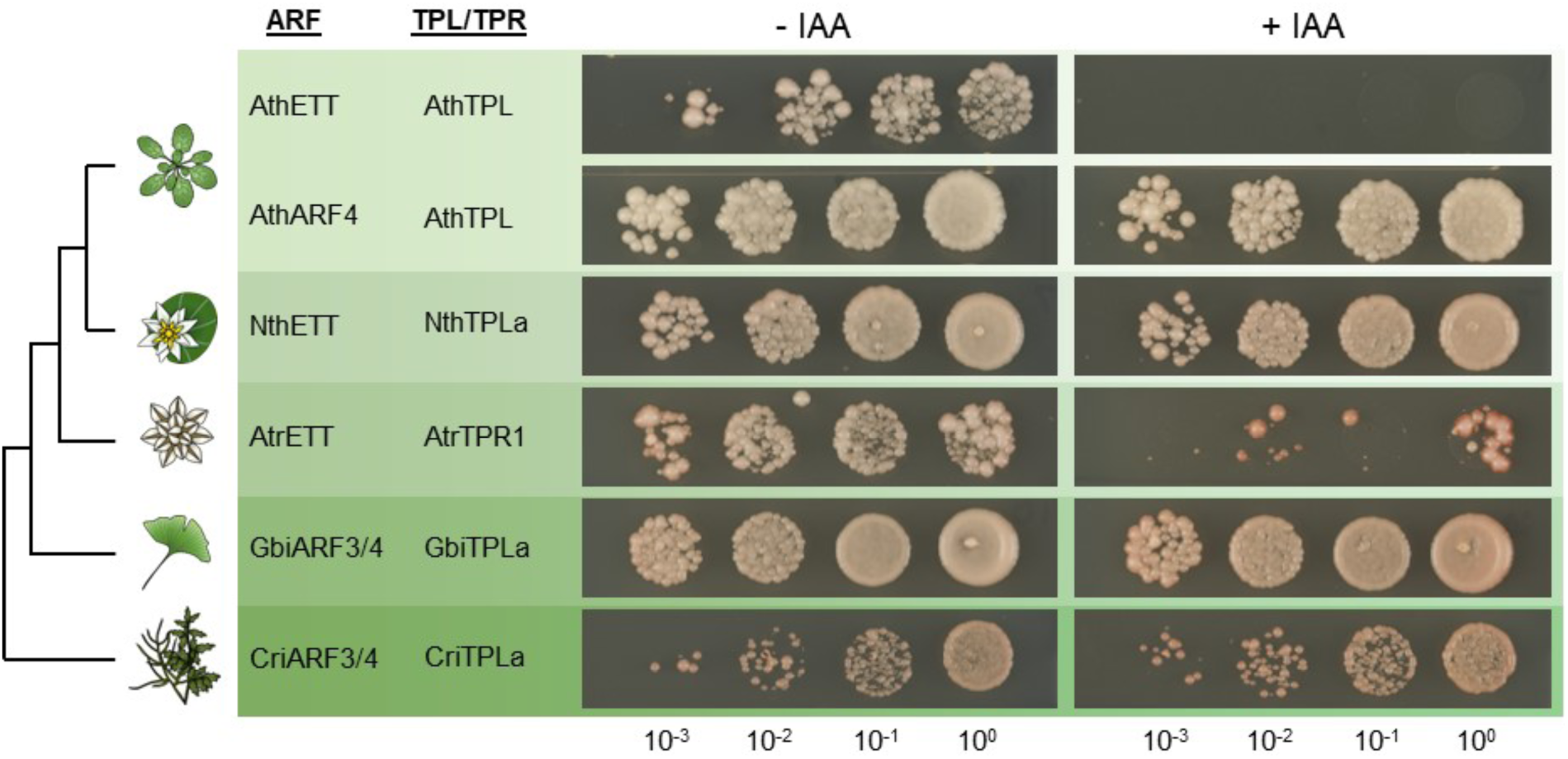
Yeast-2-hybrid auxin sensitivity screen of ARF3/4, ETT and ARF4 orthologues. Interacting ARF-TPL/TPR pairs from representative lineages were assayed for auxin sensitivity on media lacking (-IAA) or containing auxin (+IAA). Numbers at the bottom of the figure indicate the serial dilution series of the assayed yeast colonies.

In line with our previous study [31], the *A. thaliana* ETT-TPL interaction is auxin sensitive (Figure 2). In contrast, the *A. thaliana* ARF4-TPL interaction is not disrupted by auxin, indicating that auxin sensing is a property specific to ETT. The *A. trichopoda* ETT-TPR1 interaction also exhibits auxin-sensitivity while the *G. biloba* and *C. richardii* ARF3/4-TPL interactions are insensitive to auxin (Figure 2). As *A. trichopoda* belongs to a lineage that is sister to all other extant angiosperms, these results suggest that ETT-mediated auxin sensing originated after the divergence of the ETT and ARF4 clades in the last common angiosperm ancestor. However, the ETT-TPL interaction of *N. thermarum*, another ANA grade angiosperm, was not auxin sensitive (Figure 2), implying that ETT-mediated auxin signalling has been lost in this species. This result correlates with the lack of the W505 residue that have been implicated in auxin binding in the *N. thermarum* ETT orthologue [31].

### Angiosperm *ETT* orthologues fully rescue defects of the Arabidopsis *ett-3* gynoecium

In *A. thaliana*, ETT coordinates multiple aspects of gynoecium development, including floral meristem termination, valve elongation and radial style formation [25, 33, 35, 44–46]. ETT-mediated auxin signalling pathway has been shown to be important for proper establishment of the radial style [25, 31, 34]. Given that ETT, ARF4 and ARF3/4 orthologues differ in their auxin-sensing ability (Figure 2), we decided to test the ability of these orthologues to complement the gynoecium defects of the *A. thaliana ett-3* loss-of-function mutant. The sole B-ARF from the bryophyte *Marchantia polymorpha* (MpoARF2) was also included in the experiment. The coding sequence of these ARF orthologues were expressed under the control of the 5 kb *A. thaliana* ETT promoter and lines with comparable expression levels were selected for phenotyping with respect to silique length and style symmetry (measured as a medial index, Figure 3).

**Figure 3.**
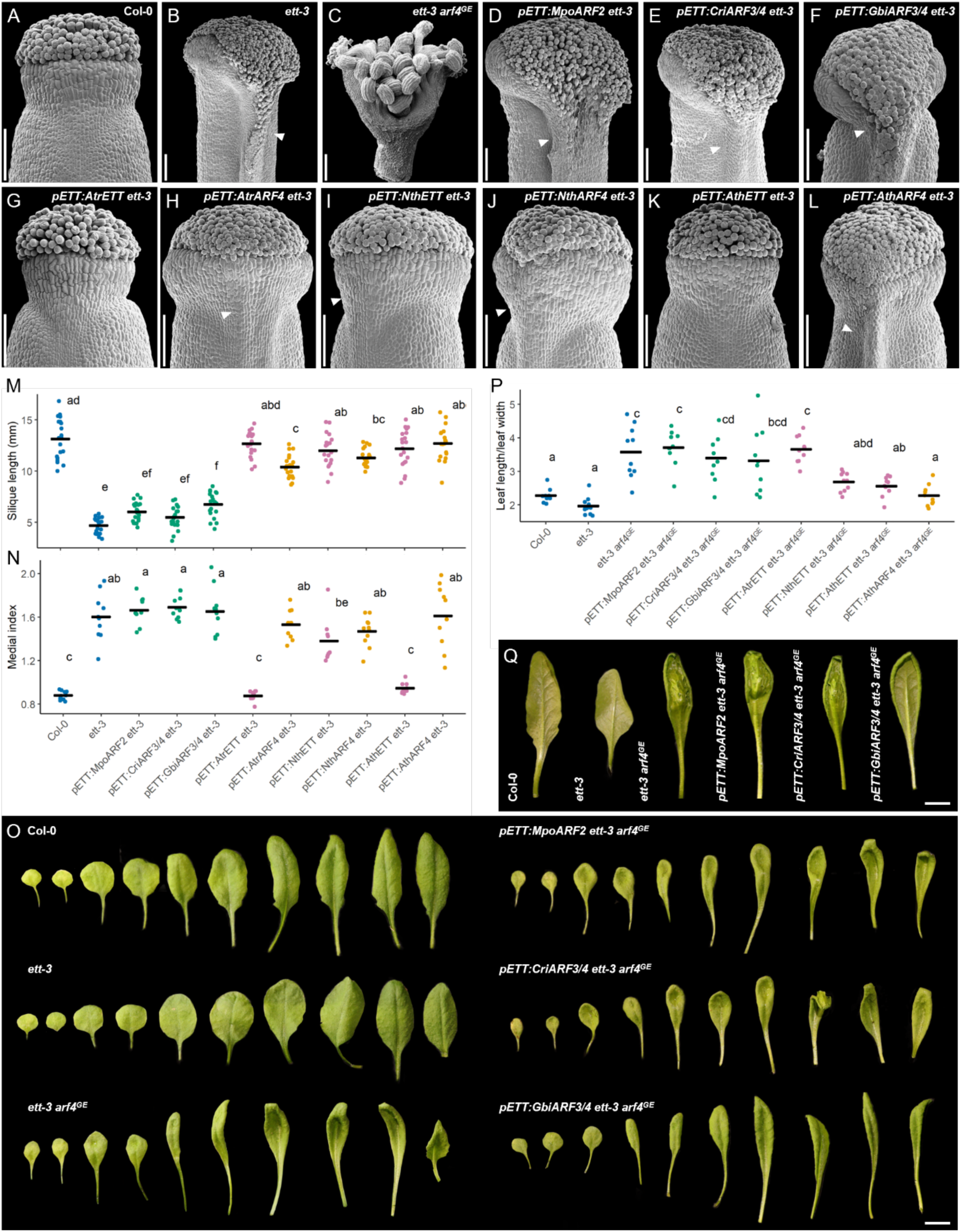
Heterologous complementation of ETT function in leaf and gynoecium development. (A-L) Gynoecium phenotypes of Col-0 (A), *ett-3* (B), and *ett-3 arf4^GE^*(C) plants and *ett-3* complementation lines expressing *MpoARF2* (D), *CriARF3/4* (E), *GbiARF3/4* (F), *AtrETT* (G)*, AtrARF4* (H)*, NthETT* (I), *NthARF4* (J)*, AthETT* (K), and *AthARF4* (L) under the control of the *AthETT* promoter. Arrows highlight medial tissue outgrowths. Scale bar, 100 µm. (M) Silique lengths of Col-0, *ett-3* and *ett-3* complementation lines (*n =* 20, black bar is the mean). (N) Medial index (ratio of maximum style width to maximum valve width) of Col-0, *ett-3* and *ett-3* complementation lines (*n =* 10, black bar is the mean). (O) Adaxial leaf phenotypes of Col-0, *ett-3,* and *ett-3 arf4^GE^* plants, and *ett-3 arf4^GE^* complementation lines expressing non-angiosperm orthologues. Scale bar, 10 mm. (P) Ratio of maximum leaf length to maximum leaf width of Col-0, *ett-3,* and *ett-3 arf4^GE^* plants, and *ett-3 arf4^GE^* complementation lines (*n =* 10, black bar is the mean). (Q) Abaxial leaf phenotypes of Col-0, *ett-3,* and *ett-3 arf4^GE^* plants, and *ett-3 arf4^GE^* complementation lines. Scale bar, 10 mm. Letters in M, N and P indicate significant differences between groups (*p* <0.05). Statistics: one-way analysis of variance followed by Tukey’s HSD post hoc test.

All angiosperm ETT orthologues were able to complement the reduced silique length phenotype of the *ett-3* mutant to levels comparable to that of the wild-type (Figure 3M). Whilst not complementing the *ett-3* gynoecium defects, all tested angiosperm ARF4 orthologues also provided substantial complementation in terms of silique growth when expressed in the *ETT* domain (Figure 3M). In contrast, silique length of non-angiosperm ARF3/4 complementation lines were comparable to that of the *ett-3* mutant, indicating that they were unable to replace AthETT function in silique growth. The *ett-3* mutant exhibits significant medial outgrowth relative to the wild-type (Figure 3A, 3B, 3N and S3). The results of the complementation experiment demonstrate that only AtrETT and AthETT were able to fully rescue the *ett-3* medial outgrowth phenotype (Figure 3G, 3K, 3M,3N and S3). Complementation lines expressing angiosperm ARF4 orthologues or the *N. thermarum* ETT still developed medial outgrowths despite the silique length complementation (Figure 3H-J, 3L, 3N and S3). Since the proteins encoding these genes are insensitive to IAA in their interaction with TPL (Figure 2), these results indicate that radial style development specifically requires the ETT-mediated non-canonical auxin signalling mechanism.

### The ETT/ARF4-like clade plays a conserved leaf polarity role in the seed plants

While ETT has a specialised role in gynoecium development, ETT and ARF4 function redundantly in the leaf adaxial-abaxial polarity and mediolateral growth network [37, 38]. This observation implies that leaf development function predates the divergence of ETT and ARF4 and was retained in both clades when ETT and ARF4 diverged in the angiosperms. To investigate this hypothesis, we first mutated the *ARF4* gene in the *ett-3* mutant background by gene editing to create the *ett-3 arf4^GE^* double mutant. In agreement with previous studies [37], this double mutant has an exacerbated gynoecium phenotype compared to the *ett-3* single mutant and is indeed sterile (Figure 3C).

In vegetative tissue, the *ett-3 arf4^GE^* mutant exhibits cup-like indentations in some young leaves, and narrow, upward curling older leaves with abaxial epidermal outgrowths (Fig. 3O-3Q and S4A). Upon introducing ETT/ARF4 orthologues under study, we found that the *M. polymorpha* ARF2 and *C. richardii* ARF3/4 orthologues were unable to rescue both these morphological defects (Figure 3O-3Q). In contrast, the *G. biloba* ARF3/4 orthologue was able to rescue the upward curling and abaxial epidermal outgrowths of older leaves (Figure 3O and 3Q) but not the reduced lamina width (Figure 3P), indicative of partial complementation. All angiosperm ETT and ARF4 orthologues were able to fully complement the *ett-3 arf4^GE^* leaf defects with an interesting novel leaf epinasty phenotype in when complementing with the *AtrETT* orthologue from Amborella (Figure 3P and S4B). These results indicate that angiosperm ETT and ARF4 orthologues, and to a lesser degree the gymnosperm ARF3/4 orthologue, were able to regulate leaf polarity and development in *A. thaliana* while non-seed plant versions were not.

### Part of the middle region domain contributes to ETT-mediated auxin sensing

Thus far, we have shown that ARF3/4, ETT and ARF4 orthologues differ in their middle region motifs and *in vivo* auxin sensitivity (Figure 1C and 2). To further investigate the contributions of the ARF domains to ETT’s novel auxin sensing ability, we performed domain swap experiments between the *G. biloba* ARF3/4, *A. trichopoda* ETT and *A. thaliana* ETT orthologues (Figure 4A). The ARF orthologues were divided into four regions: 1) the DBD, 2) MR1 encompassing the NLS and first tasiARF site, 3) MR2 encompassing the second tasiARF site, W505 residue and EAR-L motif and 4) the C-terminal PB1 domain (if present). We generated five domain swap constructs (DS1-DS5) that were tested for their auxin sensitivity in a Y2H assay with the TPL/TPR corepressors (Figure 4B).

**Figure 4.**
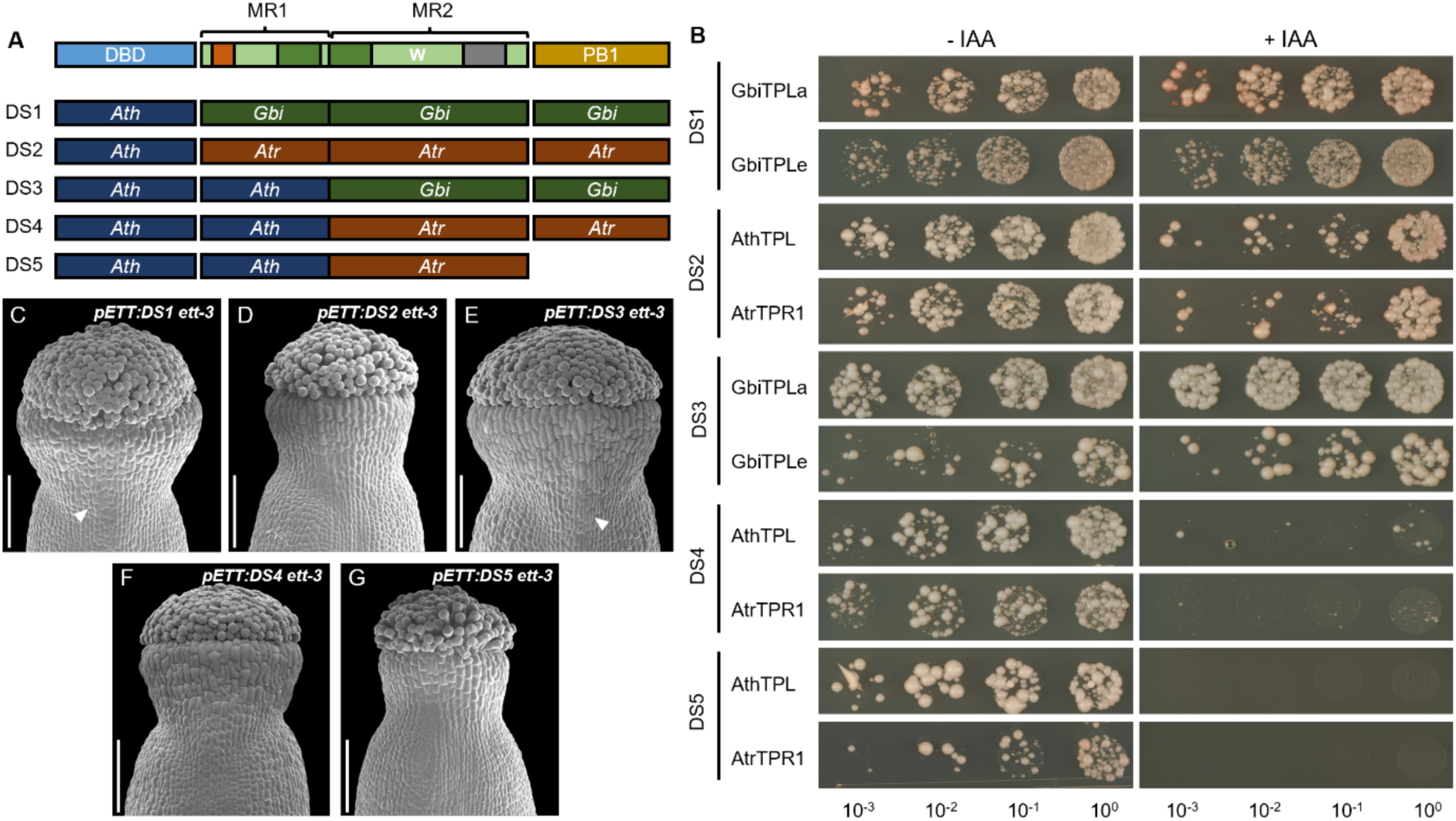
Auxin sensitivity assay and heterologous complementation ability of chimeric ETT constructs. (A) Schematic representation of the chimeric ETT constructs, DS1 to DS5. The middle region was divided into two parts (MR1 and MR2) for the domain swaps. Domains from GbiARF3/4 (*Gbi*), AtrETT (*Atr*), and AthETT (*Ath*) were used. (B) Yeast-2-hybrid auxin sensitivity screen of interactions between the chimeric ETT constructs and selected TPL/TPR partners. Yeast colonies were grown on media lacking (-IAA) and containing auxin (+IAA). Numbers at the bottom of the figure indicate the serial dilution series of the assayed yeast colonies. (C-G) Gynoecium phenotypes of *ett-3* plants expressing the chimeric ETT constructs. Arrows highlight medial tissue outgrowths. Scale bar, 100 µm.

Chimeric constructs containing the *G. biloba* ARF3/4 MR were insensitive to auxin. In contrast, all constructs expressing the *A. trichopoda* MR exhibited some degree of auxin sensitivity, including the DS4 and DS5 pair which contain or lack the *A. trichopoda* PB1 respectively. Furthermore, the difference in auxin sensitivity between the DS3 and DS4 constructs indicate that the MR2 is sufficient to influence auxin sensing. Taken together, these results suggest that MR2 is the main contributor to auxin sensing in ETT while the PB1 does not influence auxin sensing by ETT.

### Proper style morphogenesis correlates with ETT-mediated auxin sensing

As full complementation of the *ett-3* style phenotype by ETT orthologues correlate with their *in vivo* auxin sensitivity (Figure 2, 3G and 3K), we decided to test the correlation further by assessing the ability of the chimeric constructs to complement style development in the *ett-3* background (Figure 4C-G).

The DS2, DS4 and DS5 lines were able to fully complement style development in *ett-3*, whereas the DS1 and DS3 exhibited medial outgrowths similar to those from ARF4 or NthETT complementation lines (Figure 3H-J, 3L and 3N). These results indicate that auxin sensitivity of ETT in its interaction with TPL is required to overcome the style developmental defects in the *ett* mutant.

## DISCUSSION

A recurring event in land plant evolution is the diversification and co-option of ancient gene families into novel morphological contexts [47–50]. The data presented in this study provides evidence of two major functional innovations in the ARF3/4 clade that have played pivotal roles in the formation of leaves since the evolution of the gymnosperms and more recently in carpel development in angiosperms. Importantly, we show that the evolution of direct auxin sensing by ETT likely was particularly significant in the emergence of the radially symmetric style, which promotes efficient pollen tube growth and hence fertilisation in most flowering plants.

### An ancestral leaf development role for ARF3/4 in the seed plants

Our phylogenetic analysis indicates that ARF3/4 orthologues are present in the monilophytes and thus originated in the last common ancestor of the euphyllophytes (Figure 1B). The alignment of monilophyte ARF3/4 orthologues however reveal a structural divergence from the angiosperm or gymnosperm sequences, most conspicuously the lack of two tandem canonical tasiARF regulatory sites (Figure 1C and 1D). Our results, together with previously pubished data [40], indicate that monilophyte ARF3/4 orthologues are functionally divergent from seed plant orthologues. The heterologous expression of the *C. richardii* ARF3/4 orthologue failed to complement the leaf morphological defects of the *ett-3 arf4^GE^* mutant whereas partial complementation was observed for the *G. biloba* ARF3/4-expressing line (Figure 3Q). These results suggest that the leaf development role of the ARF3/4 clade emerged in the last common ancestor of the seed plants. As monilophyte fronds and seed plant leaves are considered non-homologous [51], it remains a possibility that monilophyte ARF3/4 orthologues are involved in frond development and functional studies in genetically tractable monilophyte species will be necessary to understand the role of ARF3/4 in this clade.

### Divergent evolutionary trajectories of ETT and ARF4 in the angiosperms

Previous studies revealed that ETT and ARF4 clades diverged from the ARF3/4 clade in the last common angiosperm ancestor [10, 36]. Our data indicate that ARF4 was lost twice during angiosperm radiation while the ETT clade has undergone a significant expansion in the commelinid monocots (Figure 1A and 1B). ARF4 is either partially or fully redundant with ETT in *A. thaliana* depending on tissue context while ETT paralogues in grasses have acquired novel roles in specialised floral organs [37, 52–55]. It is therefore possible that the differential patterns of ETT and ARF4 clade expansion and loss in angiosperms correlate with lineage-specific genetic redundancies and morphological specialisations.

Our data also reveal that the PB1 domain of ETT have been lost independently multiple times within monocot and magnoliid lineages (Figure 1B). The PB1 domain is functionally important for A-ARF heterodimerisation with the AUX/IAAs for regulation by the TIR1/AFB-dependent auxin signalling pathway, but its role in B-ARFs is less well understood [14, 56, 57]. While it was shown that chimeric ETT constructs carrying the *A. thaliana* ARF4 PB1 domain interfered with its function in gynoecium development [36], we do not observe any significant differences in style and silique development in complementation lines expressing full-length or truncated versions of the *A. trichopoda* ETT orthologue (Figure 3G, 3M, 3N, 4F, 4G). It is possible that the PB1 domain has no biological relevance for ETT function and thus the patterns of loss and retainment in the angiosperms reflect stochastic processes. Biochemical assays including proteomic studies with transgenic lines expressing full-length and truncated forms of ETT should be conducted to validate this hypothesis.

### Auxin sensing is an innovation of the ETT clade for style development

Our previous studies demonstrated that ETT in *A. thaliana* directly binds auxin and the TPL/TPR corepressors to control gene expression during gynoecium development [25, 31, 34]. Auxin binding by ETT is influenced by MR motifs such as the W505 residue [30, 31]. Our finding that the ETT orthologue of *A. trichopoda* also interacts with TPL/TPR proteins in an auxin-sensitive manner suggests that ETT had already gained its auxin sensing ability in the last common ancestor of the angiosperms (Figure 2). ARF3/4 or ARF4 orthologues do not exhibit auxin sensitivity in their TPL/TPR interactions, indicating that auxin sensing is specific to the ETT clade. It is thus likely that the lack of auxin sensitivity of the *N. thermarum* ETT orthologue reflects a secondary loss after the divergence of the Nymphaeales from the Amborellales. The inability of the *NthETT* to complement the *ett-3* mutant supports this. The auxin sensitivity of ETT proteins correlates with their ability to complement style symmetry establishment in the *ett-3* mutant (Figure 2, 3N, 4B-G). In contrast, all ETT and ARF4 orthologues were able to complement silique growth and flat leaf development (Figure 3M, 3P, S4). These results suggest that the ETT-mediated auxin signalling pathway might be a specialisation for style development. However, members of the ANA grade angiosperms such as *A. trichopoda* and *N. thermarum* lack conspicuous styles in their ascidiate carpels [58], so it remains unknown what the ETT-mediated auxin signalling pathway might be in *A. trichopoda*.

Our results furthermore pinpoint MR2 as the main contributing region of auxin sensing by ETT (Figure 4A and 4B). This region encompasses MR motifs that generally differ in sequence between the ETT and ARF4 clades (Figure 1C and 1D), such as the W505 residue that has been linked to auxin sensitivity [31]. However, the influence of the DBD on MR2 activity cannot be disregarded and future work focusing on whole ARF structure and motif variant activity will be important in understanding the mechanism of auxin perception by ETT.

The style is a critical structure of angiosperm fruit formation and reproduction required for efficient fertilization. Given the tight link between the auxin sensitivity of ETT and style development, it is possible that ETT has contributed to the overwhelming success of angiosperms through the non-canonical auxin signalling mechanism.

## Experimental procedures

### Plant materials and growth conditions

*Arabidopsis thaliana* ecotype Col-0 and the *ett-3* mutant allele were used in this study [25, 32]. The *arf4^GE^*line was generated using CRISPR/Cas9-mediated gene editing [59] and cross-pollinated with *ett-3* to obtain the *ett-3 arf4^GE^* double mutant. Seeds were surface sterilised, sown in petri dishes containing MS media with 0.8 % agar and 1 % sucrose, and stratified in the dark at 4 °C for three days. The seedlings were grown for 10 days under long-day conditions (16 h light/8 h dark) before they were transplanted into soil (Levington F2 compost with insecticide) under long-day conditions at 22 °C.

*Marchantia polymorpha* ecotype TAK1 and *Solanum lycopersicum* ‘MicroTom’ plants were obtained from the lab of Xiaoqi Feng (Institute of Science and Technology, Austria). *Ceratopteris richardii* ecotype Hn-n spores were obtained from Andrew Plackett (University of Birmingham) while *Nymphaea thermarum* seeds were obtained from Rebecca Povilus (Whitehead Institute) and grown at the John Innes Centre. *Amborella trichopoda* tissue were collected from Cambridge University Botanic Gardens, while *Ginkgo biloba* tissue was collected from a tree in the John Innes Centre courtyard.

### Phylogeny construction and motif identification

A total of 104 proteomes from sequenced genomes available mainly through PLAZA [60] and Phytozome [61] were searched for ARF homologs using HMMER v3.3 [62] with HMM made from the alignment of Arabidopsis and Marchantia ARF homologs as query sequences. The initial fast alignment was made using the MAFFT FFT-NS-1 algorithm v7.505 [63] and a tree was built with IQtree v2.2.0 [64] with a maximum of 100 rapid bootstraps. From this tree, only the sequences in the B-ARF clade were selected along with a few A-ARF sequences that were used as outgroups for the later phylogenetic tree construction. These selected B-ARF sequences were further aligned using MAFFT E-INS-i algorithm v7.505 [63]. Regions with more than 60% gaps were removed from the alignment using trimAl [65]. IQtree2 was used with JTT+F+R10 as the evolutionary model selected from ModelFinder [66] with 1000 rapid bootstraps. Phylogenetic trees were visualised using iTOL v5 [67].

From our alignment, the ARF MR was selected in JalView v2 [68] using sequences of the DNA-binding domain [69] and the PB1 domain [70]. The R packages ‘protr’ and ‘Peptides’ were used to compute sequence features of the MR [71, 72]. Sequence logos for visualising conserved peptides were made using WebLogo v3.7.4 [73].

### Plasmid construction

The RNeasy Plant Mini Kit (Qiagen) was used to isolate RNA from *A. thaliana* inflorescences and shoots. For all other species, a CTAB pre-treatment protocol was utilised [74]. The on-column RNAse free DNAse kit (Qiagen) was added to remove genomic DNA and the SuperScript™ IV First-Strand Synthesis kit (ThermoFisher) was used according to synthesise cDNA from 1 μg of total RNA. The coding regions of ARF and TPL/TPR orthologues were cloned from cDNA using the Phusion Flash High-Fidelity PCR Master Mix kit (Thermo Scientific). The list of primers used are shown in Table S3.

In-Fusion cloning (Takara Bio) was used for the generation of all constructs in this study. For the Y2H constructs, the coding sequence of the orthologues were inserted into linearised pB1880 Gal4-BD and pB1881 Gal4-AD vectors [75] while the 5 kb *A. thaliana ETT* promoter and ARF orthologue coding sequence were inserted into the pCambia1305 vector for the *in planta* complementation constructs [75]. Plasmids were extracted using the NucleoSpin Plasmid EasyPure kit (Macherey-Nagel) and sequences were validated via Sanger sequencing with Macrogen Europe.

### Generation and phenotyping of transgenic plants

The constructs were introduced into *Agrobacterium tumefaciens* strain GV3101 via electroporation for plant transformation using the floral dip method [76]. Transformed *ett-3* plants were selected on MS plates containing 30 μg/ml hygromycin and transplanted into soil after 10 days. Plants were genotyped in the T1 generation for ARF transgene presence. Single copy T-DNA insertion lines were selected in the T2 generation and homozygous plants were selected for in the T3 generation through assessment of transgene segregation ratios. Quantitative PCR was used to select plant lines of comparable transgene expression levels.

To obtain the *ett-3 arf4^GE^* complementation lines, selected *ett-3* complementation lines were cross-pollinated with *ett-3* (-/-) *arf4^GE^* (+/-) plants. The presence of transgenes and the homozygous *arf4^GE^* mutant allele was checked through genotyping and segregation analyses in the F2 generation.

To calculate the medial index, the maximum style width to style height ratio was measured and calculated from SEM images of gynoecia using the ImageJ software. Fully elongated but unripe siliques were collected, lined on a piece of paper, and measured for silique lengths.

The seventh to ninth leaves of *A. thaliana* rosettes were collected, laid flat and photographed for leaf shape quantification. The maximum width and length of the lamina were measured in ImageJ and the ratio of leaf length to leaf width for each line was calculated.

### Scanning electron microscopy

Inflorescences were collected in vials and fixed in FAA solution (3.7 % formaldehyde, 3 % acetic acid and 50 % ethanol) at room temperature overnight. After the removal of the fixative, 70 % ethanol was added to the vials and left to incubate overnight. A dehydration series consisting of 90 % ethanol for 1 hour, 100 % ethanol for 1 hour and two washes of 100 % dry ethanol for 30 min each was used to prepare the samples prior to critical point drying using the Leica CPD300. The dried gynoecia were dissected and mounted on stubs for gold coating using an Agar high resolution sputter coater. The Zeiss Supra 55VP Field Emission Scanning Electron Microscope (3kV acceleration voltage) was used to image the samples.

### Yeast-2-hybrid assays

The AH109 yeast strain (Clontech) was used for all Y2H experiments. Yeast cells were transformed using the co-transformation method [77]. Yeast cells were grown at 28 °C for 3-4 days on WL YSD minimal media lacking tryptophan and leucine to select for positive transformants. The transformed yeast colonies were then plated on WLAH media lacking tryptophan, leucine, adenine, and histidine to assay for protein-protein interactions. To test for auxin sensitivity, yeast cells were serially diluted (100, 10-1, 10-2 and 10-3) and spotted on WLAH media supplemented with 100 μM indole-3-acetic acid (IAA). Images were taken after 5 days of growth at 28 °C.

### Quantification and Statistical Analysis

All statistical analyses and data visualisation were performed in RStudio. One-way ANOVA followed by Tukey’s HSD post hoc test was used to compare differences between plant lines for medial index, silique length and leaf shape quantifications. In all cases, *P* values less than 0.05 were considered significant.

## ACKNOWLEDGMENTS

We thank Yuli Ding and Yuelin Zhang (University of British Columbia) for sharing the pCambia1305, pB1880/pB1881 and CRISPR plasmids, and André Kuhn (Wageningen University) and Yuxiang Jiang (Leiden University) for help with the Y2H experiments and *ett-3 arf4^GE^* crosses. We are grateful to James Walker (Salk Institute) for providing *M. polymorpha* TAK1 plants, Andy Plackett (University of Birmingham) for *C. richardii* Hnn spores, Rebecca Povilus and William Friedman (Arnold Arboretum, Harvard University) for *N. thermarum* seeds, and Cambridge University Botanic Garden for *A. trichopoda* samples. We also thank Norwich Research Park Bioimaging for assistance with the SEM. This study was supported by a grant from the Biotechnological and Biological Research Council (BBSRC) to L.Ø. (BB/P020747/1) and an Institute Strategic Programme Grant from the BBSRC to the John Innes Centre (BB/P013511/1). A.C.H.A. was supported by the Sainsbury PhD studentship.

## AUTHOR CONTRIBUTIONS

A.C.H.A. and L. Ø. designed the study. A.C.H.A. conducted the majority of the experiments. S.M. and E.V.D.L. conducted the phylogenetic and bioinformatic analyses. L. Ø. and D. W. supervised the study. A.C.H.A. wrote the manuscript with revisions from L. Ø. All authors commented and agreed on the manuscript before submission.

