## Supplemental Information for "ETTIN-mediated auxin signalling is an angiosperm-specific neofunctionalization for carpel development"

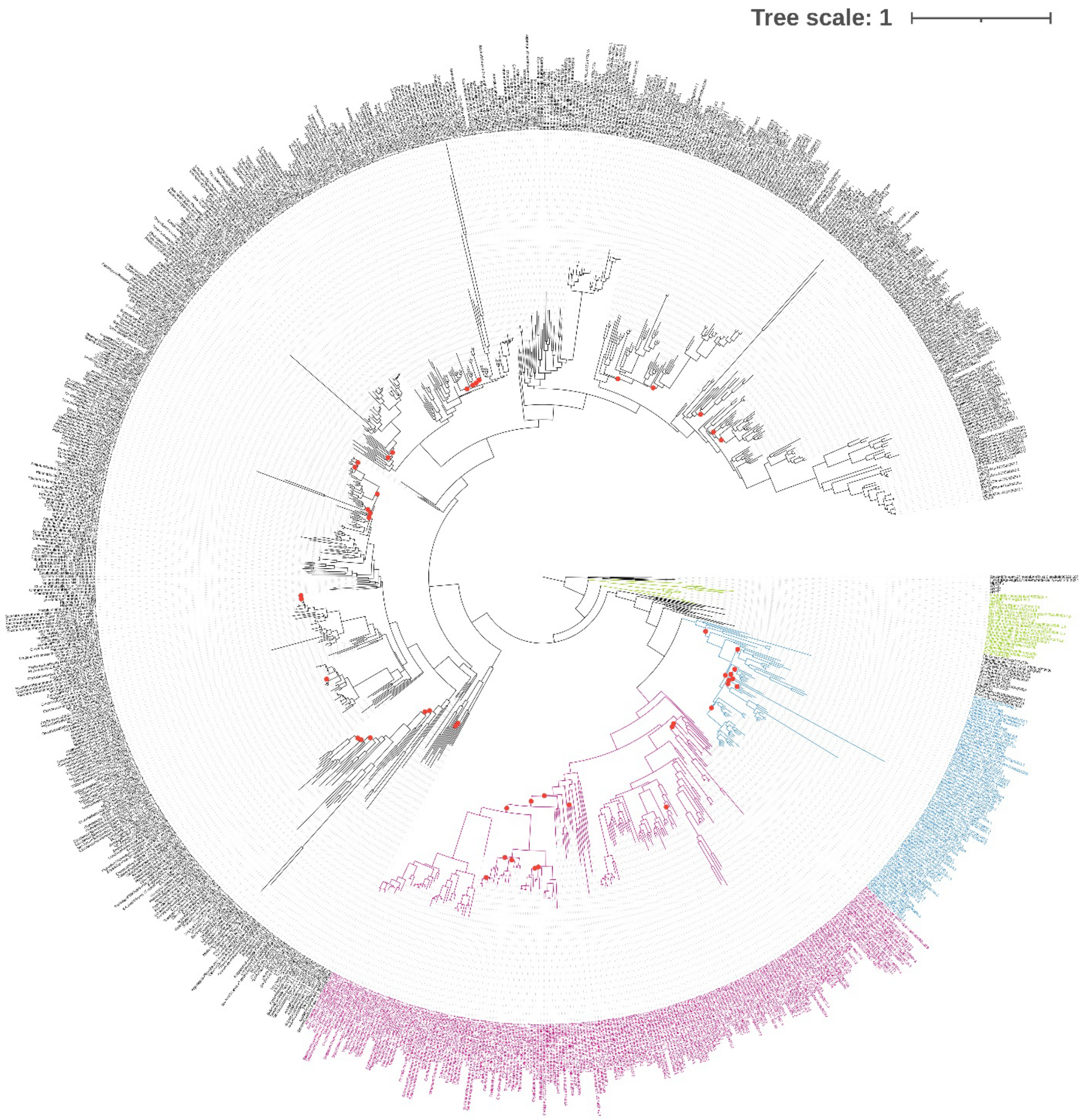


**Figure S1: Full B-ARF phylogeny of the land plants.** The colour-coded segment on the lower right is identical to Figure 1A. The ARF2 and ARF1/rest clades are represented by the black segment on the upper left side. Red circles on the branches indicate poor bootstrap support (<75) for that split.

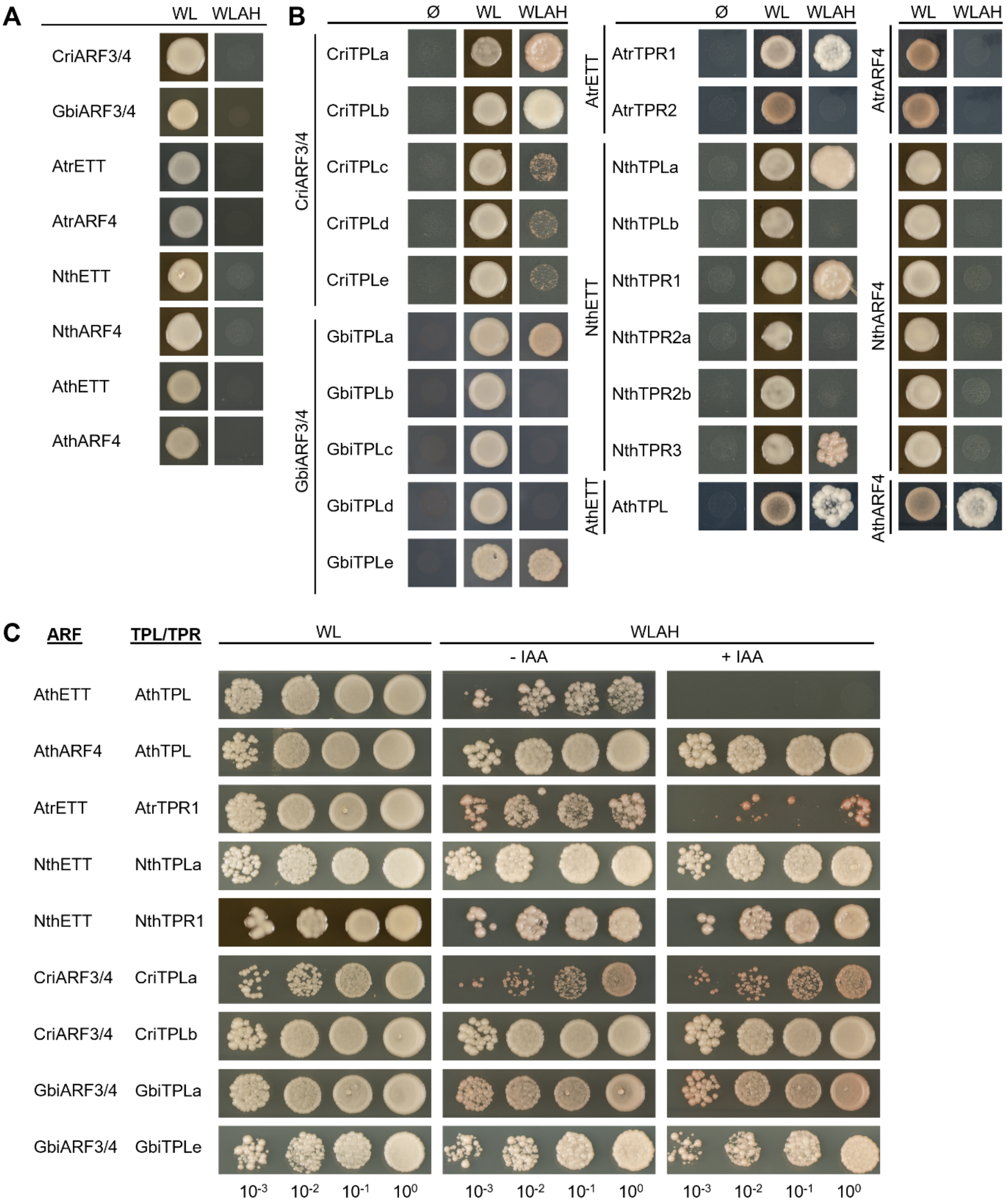


**Figure S2: Yeast-2-hybrid screen of positive ARF-TPL/TPR interactions across the land plants.**
(A) Auto-activation test for ARF Y2H constructs.
(B) Yeast-2-hybrid screen showing positive or negative interactions between ARF and TPL/TPR pairs. Ø represents the empty vector control to test for auto-activation of TPL/TPR constructs. WL indicates growth of colonies on media lacking tryptophan and leucine. WLAH indicates growth of colonies on media lacking tryptophan, leucine, adenine and histidine.
(C) Auxin sensitivity assay of positive ARF-TPL/TPR interactions. Colony growth was assessed on media lacking auxin (- IAA) or containing auxin (+ IAA). Numbers at the bottom of the figure indicate the serial dilution series of the assayed yeast colonies.

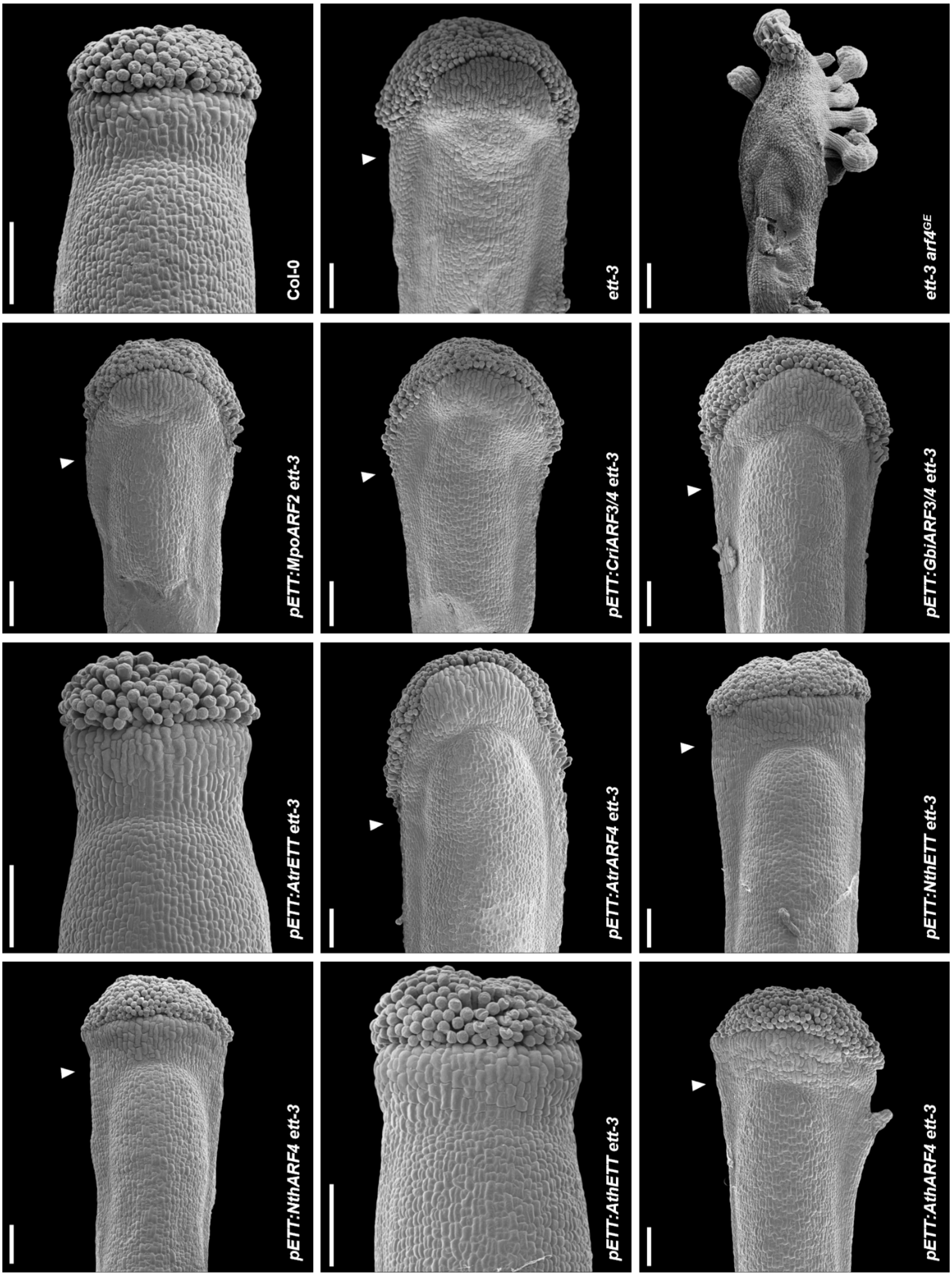


**Figure S3: Lateral view of the gynoecium of wild-type, mutant and complementation lines.** Arrows highlight medial tissue outgrowths. Scale bar, 100 µm.

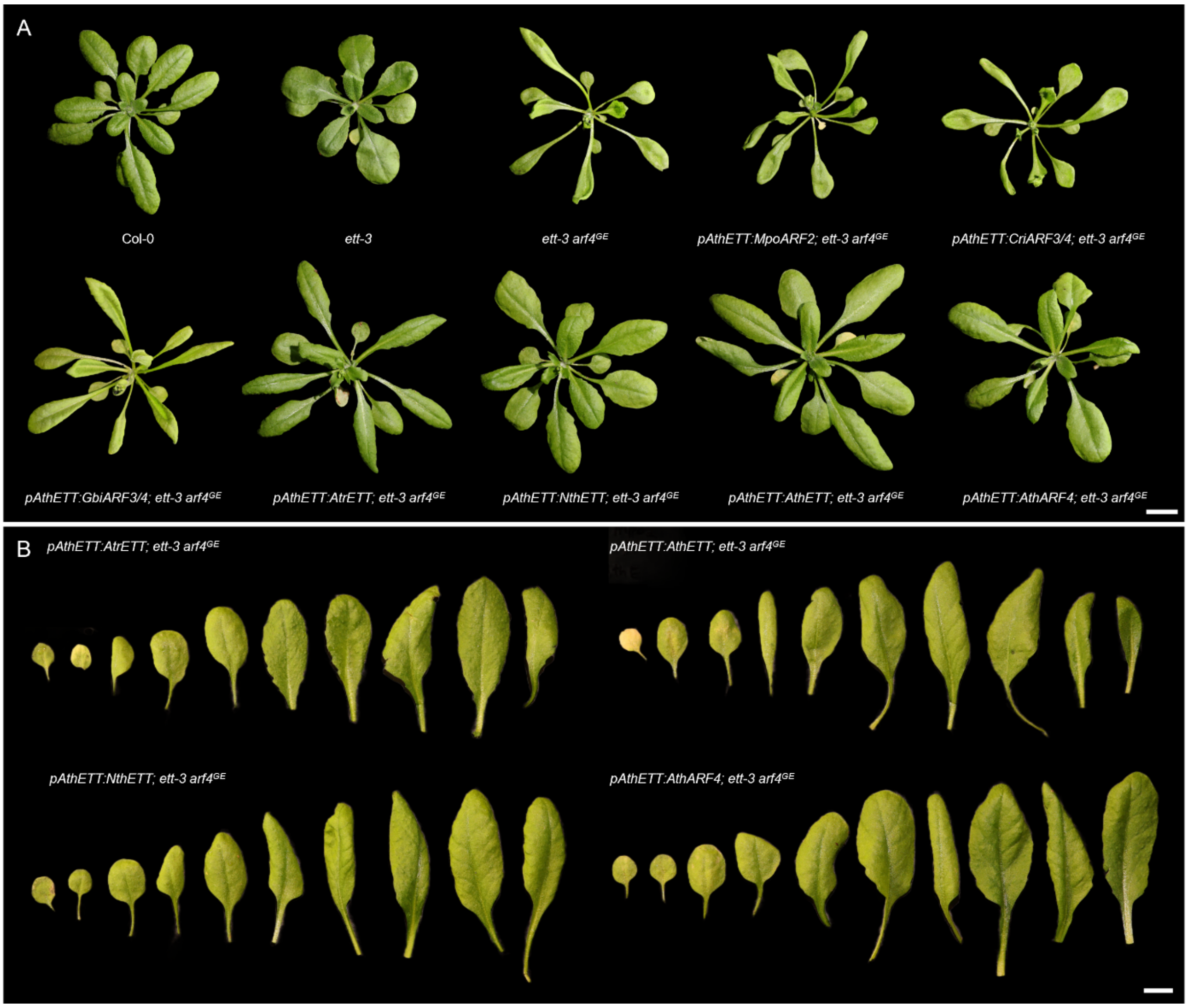


**Figure S4: Vegetative phenotypes of control and *ett-3 arf4$^{GE}$* complementation lines.**
(A) Rosette phenotypes of Col-0, *ett-3,* and *ett-3 arf4$^{GE}$* plants, and *ett-3 arf4$^{GE}$* complementation lines. Scale bar, 10 mm.
(B) Adaxial leaf series of *ett-3 arf4$^{GE}$* complementation lines expressing angiosperm ETT or ARF4 constructs. Scale bar, 10 mm.

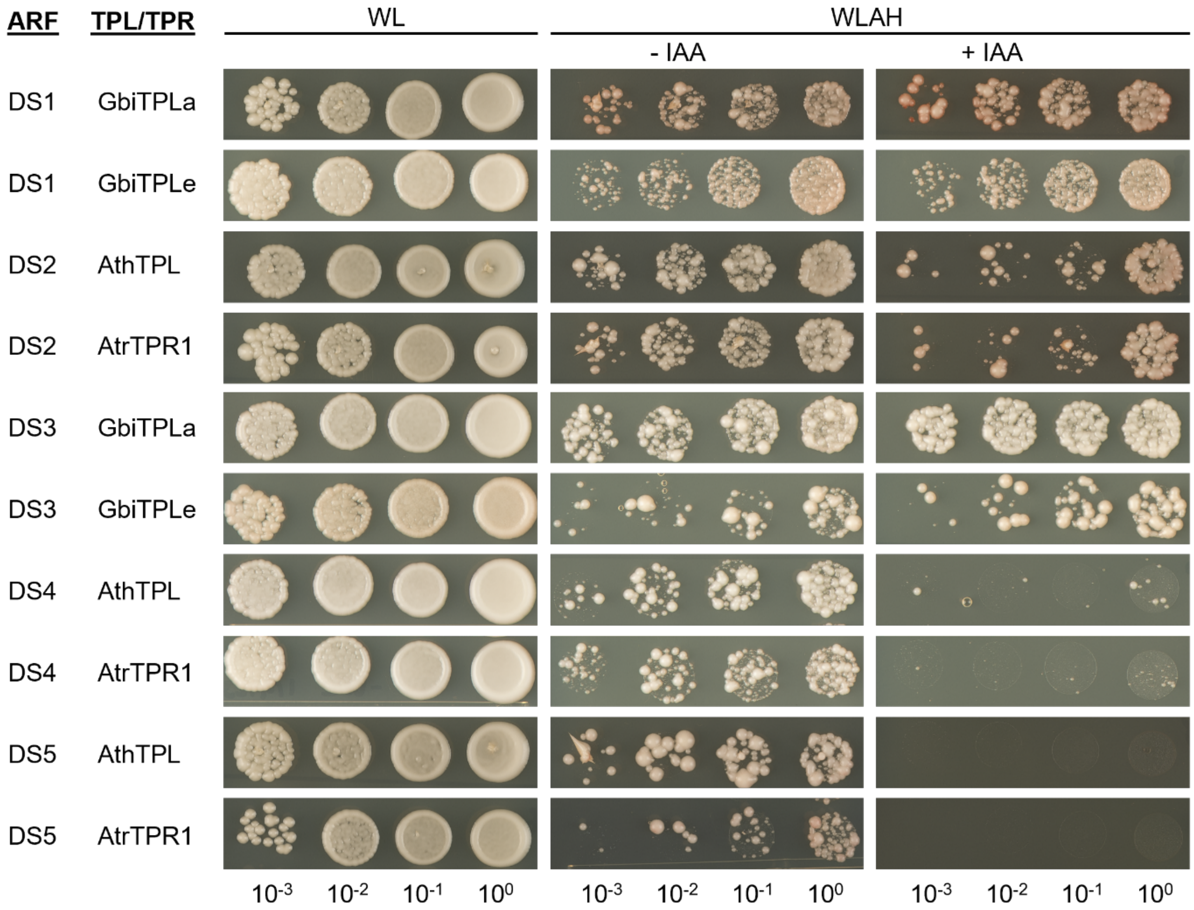


**Figure S5: Auxin sensitivity yeast-2-hybrid assay of chimeric ARF-TPL/TPR interactions.** DS1-DS5 were defined in Figure 4A. WL indicates growth of colonies on media lacking tryptophan and leucine. WLAH indicates growth of colonies on media lacking tryptophan, leucine, adenine and histidine. Colony growth was assessed on media lacking auxin (- IAA) or containing auxin (+ IAA). Numbers at the bottom of the figure indicate the serial dilution series of the assayed yeast colonies.

**Table S1: Genomes included in the B-ARF phylogeny.** Manually sourced genomes were downloaded from links provided in the referenced publication.

| **Species** | **Clade** | **Source** | **Reference** |
|---|---|---|---|
| *Chlamydomonas reinhardtii* | Alga: Chlorophyte | PLAZA | Merchant et al. (2007) |
| *Micromonas commoda* | Alga: Chlorophyte | PLAZA | Worden et al. (2009) |
| *Mesotaenium endlicherianum* | Alga: Charophyte | Manual | Cheng et al. (2019) |
| *Chara braunii* | Alga: Charophyte | Manual | Nishiyama et al. (2018) |
| *Mesostigma viride* | Alga: Charophyte | Manual | Wang et al. (2020) |
| *Chlorokybus atmophyticus* | Alga: Charophyte | Manual | Wang et al. (2020) |
| *Coleochaete scutata* | Alga: Charophyte | Manual | de Vries et al. (2018) |
| *Klebsormidium nitens* | Alga: Charophyte | Manual | Hori et al. (2014) |
| *Spirogloea muscicola* | Alga: Charophyte | Manual | Cheng et al. (2019) |
| *Penium margaritaceum* | Alga: Charophyte | Manual | Jiao et al. (2020) |
| *Marchantia palacea* | Bryophyte: Liverwort | Manual | Diop et al. (2020) |
| *Marchantia polymorpha* | Bryophyte: Liverwort | Manual | Bowman et al. (2017) |
| *Physcomitrium patens* | Bryophyte: Moss | Manual | Lang et al. (2018) |
| *Ceratodon purpureus* | Bryophyte: Moss | Phytozome | Carey et al. (2021) |
| *Sphagnum fallax* | Bryophyte: Moss | Phytozome | Healey et al. (2023) |
| *Sphagnum magellanicum* | Bryophyte: Moss | Phytozome | Healey et al. (2023) |
| *Anthoceros agrestis* | Bryophyte: Hornwort | Manual | Li et al. (2020b) |
| *Selaginella moellendorffii* | Lycophyte | Manual | Banks et al. (2011) |
| *Azolla filiculoides* | Monilophyte | Manual | Li et al. (2018) |
| *Salvinia cucullata* | Monilophyte | Manual | Li et al. (2018) |
| *Ceratopteris richardii* | Monilophyte | Manual | Marchant et al. (2022) |
| *Ginkgo biloba* | Gymnosperm | Manual | Liu et al. (2021) |
| *Pinus taeda* | Gymnosperm | Manual | Zimin et al. (2017) |
| *Picea abies* | Gymnosperm | Manual | Nystedt et al. (2013) |
| *Gnetum montanum* | Gymnosperm | Manual | Wan et al. (2018) |
| *Amborella trichopoda* | Angiosperm: ANA Grade | Manual | Project et al. (2013) |
| *Nymphaea thermarum* | Angiosperm: ANA Grade | Manual | Povilus et al. (2020) |
| *Aristolochia fimbriata* | Angiosperm: Magnoliid | Manual | Qin et al. (2021) |

| | | | |
|---|---|---|---|
| *Piper nigrum* | Angiosperm: Magnoliid | Manual | Hu et al. (2019) |
| *Liriodendron chinense* | Angiosperm: Magnoliid | Manual | Chen et al. (2019) |
| *Chimonanthus salicifolius* | Angiosperm: Magnoliid | Manual | Lv et al. (2020) |
| *Persea americana* | Angiosperm: Magnoliid | Manual | Rendón-Anaya et al. (2019) |
| *Cinnamomum kanehirae* | Angiosperm: Magnoliid | Phytozome | Chaw et al. (2019) |
| *Zostera marina* | Angiosperm: Monocot | PLAZA | Olsen et al. (2016) |
| *Spirodela polyrhiza* | Angiosperm: Monocot | PLAZA | Wang et al. (2014b) |
| *Apostasia shenzhenica* | Angiosperm: Monocot | PLAZA | Zhang et al. (2017) |
| *Phalaenopsis equestris* | Angiosperm: Monocot | PLAZA | Cai et al. (2015) |
| *Asparagus offinalis* | Angiosperm: Monocot | PLAZA | Harkess et al. (2017) |
| *Elaeis guineensis* | Angiosperm: Monocot | PLAZA | Singh et al. (2013) |
| *Calamus simplicifolius* | Angiosperm: Monocot | PLAZA | Zhao et al. (2018b) |
| *Musa acuminata* | Angiosperm: Monocot | PLAZA | D'Hont et al. (2012) |
| *Ananas comosus* | Angiosperm: Monocot | PLAZA | Ming et al. (2015) |
| *Zoysia japonica* | Angiosperm: Monocot | PLAZA | Tanaka et al. (2016) |
| *Oropetium thomaeum* | Angiosperm: Monocot | PLAZA | VanBuren et al. (2018) |
| *Brachypodium distachyon* | Angiosperm: Monocot | Manual | Vogel et al. (2010) |
| *Triticum turgidum* | Angiosperm: Monocot | PLAZA | Maccaferri et al. (2019) |
| *Triticum aestivum* | Angiosperm: Monocot | PLAZA | Consortium et al. (2014) |
| *Lolium perenne* | Angiosperm: Monocot | PLAZA | Nagy et al. (2022) |
| *Hordeum vulgare* | Angiosperm: Monocot | PLAZA | Mascher et al. (2017) |
| *Phyllostachys edulis* | Angiosperm: Monocot | PLAZA | Zhao et al. (2018a) |
| *Saccharum spontaneum* | Angiosperm: Monocot | PLAZA | Zhang et al. (2018a) |
| *Zea mays* | Angiosperm: Monocot | Manual | Schnable et al. (2009) |
| *Zea mays* PH207 | Angiosperm: Monocot | PLAZA | Hirsch et al. (2016) |
| *Setaria italica* | Angiosperm: Monocot | PLAZA | He et al. (2023) |
| *Sorghum bicolor* | Angiosperm: Monocot | PLAZA | Paterson et al. (2009) |
| *Cenchrus americanus* | Angiosperm: Monocot | PLAZA | Varshney et al. (2017) |
| *Oryza brachyantha* | Angiosperm: Monocot | PLAZA | Chen et al. (2013) |
| *Oryza sativa ssp. indica* | Angiosperm: Monocot | PLAZA | Yu et al. (2002) |

| | | | |
|---|---|---|---|
| *Oryza sativa ssp. japonica* | Angiosperm: Monocot | Manual | Sasaki and International Rice Genome Sequencing (2005) |
| *Nelumbo nucifera* | Angiosperm: Eudicot | PLAZA | Ming et al. (2013) |
| *Vitis vinifera* | Angiosperm: Eudicot | Manual | Jaillon et al. (2007) |
| *Pyrus bretschneideri* | Angiosperm: Eudicot | PLAZA | Wu et al. (2013) |
| *Malus domestica* | Angiosperm: Eudicot | PLAZA | Daccord et al. (2017) |
| *Fragaria vesca* | Angiosperm: Eudicot | PLAZA | Shulaev et al. (2011) |
| *Prunus persica* | Angiosperm: Eudicot | PLAZA | Verde et al. (2013) |
| *Ziziphus jujuba* | Angiosperm: Eudicot | PLAZA | Liu et al. (2014a) |
| *Citrullus lanatus* | Angiosperm: Eudicot | PLAZA | Guo et al. (2013) |
| *Cucumis melo* | Angiosperm: Eudicot | PLAZA | Garcia-Mas et al. (2012) |
| *Cucumis sativus* | Angiosperm: Eudicot | PLAZA | Hu et al. (2019) |
| *Trifolium pratense* | Angiosperm: Eudicot | PLAZA | De Vega et al. (2015) |
| *Arachis ipaensis* | Angiosperm: Eudicot | PLAZA | Lu et al. (2018) |
| *Cajanus cajan* | Angiosperm: Eudicot | PLAZA | Varshney et al. (2012) |
| *Vigna radiata* | Angiosperm: Eudicot | PLAZA | Kang et al. (2014) |
| *Medicago truncatula* | Angiosperm: Eudicot | PLAZA | Young et al. (2011) |
| *Cicer arietinum* | Angiosperm: Eudicot | PLAZA | Varshney et al. (2013) |
| *Glycine max* | Angiosperm: Eudicot | PLAZA | Schmutz et al. (2010) |
| *Populus trichocarpa* | Angiosperm: Eudicot | PLAZA | Tuskan et al. (2006) |
| *Ricinus communis* | Angiosperm: Eudicot | PLAZA | (Chan et al., 2010) |
| *Manihot esculenta* | Angiosperm: Eudicot | PLAZA | Wang et al. (2014a) |
| *Hevea brasiliensis* | Angiosperm: Eudicot | PLAZA | Tang et al. (2016) |
| *Citrus clementina* | Angiosperm: Eudicot | PLAZA | Wu et al. (2014) |
| *Eucalyptus grandis* | Angiosperm: Eudicot | PLAZA | Myburg et al. (2014) |
| *Gossypium raimondii* | Angiosperm: Eudicot | PLAZA | Wang et al. (2012) |
| *Corchorus olitorius* | Angiosperm: Eudicot | PLAZA | Islam et al. (2017) |
| *Theobroma cacao* | Angiosperm: Eudicot | PLAZA | Argout et al. (2011) |
| *Carica papaya* | Angiosperm: Eudicot | PLAZA | Ming et al. (2008) |
| *Tarenaya hassleriana* | Angiosperm: Eudicot | PLAZA | Cheng et al. (2013) |
| *Brassica rapa* | Angiosperm: Eudicot | PLAZA | Wang et al. (2011) |

| | | | |
|---|---|---|---|
| *Brassica oleracea* | Angiosperm: Eudicot | PLAZA | Liu et al. (2014b) |
| *Schrenkiella parvula* | Angiosperm: Eudicot | PLAZA | Dassanayake et al. (2011) |
| *Capsella rubella* | Angiosperm: Eudicot | PLAZA | Slotte et al. (2013) |
| *Arabidopsis lyrata* | Angiosperm: Eudicot | PLAZA | Hu et al. (2011) |
| *Arabidopsis thaliana* | Angiosperm: Eudicot | PLAZA | The Arabidopsis Genome (2000) |
| *Beta vulgaris* | Angiosperm: Eudicot | PLAZA | Dohm et al. (2014) |
| *Chenopodium quinoa* | Angiosperm: Eudicot | PLAZA | Jarvis et al. (2017) |
| *Actinidia chinensis* | Angiosperm: Eudicot | PLAZA | Huang et al. (2013a) |
| *Daucus carota* | Angiosperm: Eudicot | PLAZA | Iorizzo et al. (2016) |
| *Erythranthe guttata* | Angiosperm: Eudicot | PLAZA | Hellsten et al. (2013) |
| *Utricularia gibba* | Angiosperm: Eudicot | PLAZA | Ibarra-Laclette et al. (2013) |
| *Coffea canephora* | Angiosperm: Eudicot | PLAZA | Denoeud et al. (2014) |
| *Capsicum annuum* | Angiosperm: Eudicot | PLAZA | Kim et al. (2014) |
| *Solanum lycopersicum* | Angiosperm: Eudicot | Manual | Sato et al. (2012) |
| *Solanum tuberosum* | Angiosperm: Eudicot | PLAZA | Xu et al. (2011) |
| *Petunia axillaris* | Angiosperm: Eudicot | PLAZA | Bombarely et al. (2016) |

**Table S2:** List of ARF and TPL/TPR orthologues used in the Y2H screens.

| Species | Gene | ID | Reference |
|---|---|---|---|
| *Ceratopteris richardii* | *CriARF3/4* | Ceric.21G089900 | Marchant et al. (2022); Phytozome |
| | *CriTPLa* | Ceric.08G073000 | |
| | *CriTPLb* | Ceric.07G013800 | |
| | *CriTPLc* | Ceric.1Z260400 | |
| | *CriTPLd* | Ceric.14G074400 | |
| | *CriTPLe* | Ceric.10G065300 | |
| *Ginkgo biloba* | *GbiARF3/4* | Gb_17830/Gb_17831 | Guan et al. (2016); Liu et al. (2021) |
| | *GbiTPLa* | Gb_37153 | |
| | *GbiTPLb* | Gb_32130 | |
| | *GbiTPLc* | Gb_04053 | |
| | *GbiTPLd* | Gb_26928 | |
| | *GbiTPLe* | Gb_39002 | |
| *Amborella trichopoda* | *AtrETT* | AMTR_s00021p00200760 | Project et al. (2013) |
| | *AtrARF4* | AMTR_s00034p00110140 | |
| | *AtrTPR1* | AMTR_s00051p00079490 | |
| | *AtrTPR2* | AMTR_s00048p00159380 | |
| *Nymphaea thermarum* | *NthETT* | EJ110_NYTH15956 | Povilus et al. (2020) |
| | *NthARF4* | EJ110_NYTH47885 | |
| | *NthTPLa* | EJ110_NYTH22400 | |
| | *NthTPLb* | EJ110_NYTH18194 | |
| | *NthTPR1* | EJ110_NYTH02857 | |
| | *NthTPR2a* | EJ110_NYTH44705 | |
| | *NthTPR2b* | EJ110_NYTH49689 | |
| | *NthTPR3* | EJ110_NYTH09053 | |
| *Arabidopsis thaliana* | *AthETT* | AT2G33860 | The Arabidopsis Genome (2000); TAIR |
| | *AthARF4* | AT5G60450 | |
| | *AthTPL* | AT1G15750 | |

**Table S3.** List of oligonucleotides used in this study.

| Orientation | Gene | Sequence | Use |
|---|---|---|---|
| Forward | *CriARF3/4* | ACGACAAGGGGTCGACCATGCATCTCGACTCGCAGC | Y2H: pB1880/81 |
| Reverse | *CriARF3/4* | ACCGCGGTGGCGGCCGCTTAACCTGAGATGGAGTTCCTG | Y2H: pB1880 |
| Reverse | *CriARF3/4* | ACTTACTTAGCGGCCGCTTAACCTGAGATGGAGTTCCTG | Y2H: pB1881 |
| Forward | *GbiARF3/4* | ACGACAAGGGGTCGACCATGGAAATTGATCTCAACAGTCC | Y2H: pB1880/81 |
| Reverse | *GbiARF3/4* | ACCGCGGTGGCGGCCGCTTAAATTCTTGTTGCTGGTGGAG | Y2H: pB1880 |
| Reverse | *GbiARF3/4* | ACTTACTTAGCGGCCGCTTAAATTCTTGTTGCTGGTGGAG | Y2H: pB1881 |
| Forward | *AtrETT* | ACGACAAGGGGTCGACCATGGGCATTGATCTGAACCGG | Y2H: pB1880/81 |
| Reverse | *AtrETT* | ACCGCGGTGGCGGCCGCTTACACAGCTCTAGCAAGGCC | Y2H: pB1880 |
| Reverse | *AtrETT* | ACTTACTTAGCGGCCGCTTACACAGCTCTAGCAAGGCC | Y2H: pB1881 |
| Forward | *AtrARF4* | ACGACAAGGGGTCGACCATGGAAATTGATCTCAACTGCG | Y2H: pB1880/81 |
| Reverse | *AtrARF4* | ACCGCGGTGGCGGCCGCCTATTCGAGTCCTCTTGTTATGG | Y2H: pB1880 |
| Reverse | *AtrARF4* | ACTTACTTAGCGGCCGCCTATTCGAGTCCTCTTGTTATGG | Y2H: pB1881 |
| Forward | *NthETT* | ACGACGACAAGGGGTCGACCATGGAGATCGATCTAAACAGGG | Y2H: pB1880/81 |
| Reverse | *NthETT* | ACCGCGGTGGCGGCCGCTCAAGTCCTTGTTACCGTCAC | Y2H: pB1880 |
| Reverse | *NthETT* | ACTTACTTAGCGGCCGCTCAAGTCCTTGTTACCGTCAC | Y2H: pB1881 |
| Forward | *NthARF4* | ACGACAAGGGGTCGACCATGGAAATTGATCTCAACAGTGCAC | Y2H: pB1880/81 |
| Reverse | *NthARF4* | ACCGCGGTGGCGGCCGCCTAGATGTTGTTTTCCTTAGTCAAC | Y2H: pB1880 |
| Reverse | *NthARF4* | ACTTACTTAGCGGCCGCCTAGATGTTGTTTTCCTTAGTCAAC | Y2H: pB1881 |
| Forward | *AthETT* | ACGACAAGGGGTCGACCATGGGTGGTTTAATCGATCTGAACG | Y2H: pB1880/81 |
| Reverse | *AthETT* | ACCGCGGTGGCGGCCGCCTAGAGAGCAATGTCTAGCAACATG | Y2H: pB1880 |
| Reverse | *AthETT* | ACTTACTTAGCGGCCGCCTAGAGAGCAATGTCTAGCAACATG | Y2H: pB1881 |
| Forward | *AthARF4* | ACGACAAGGGGTCGACCATGGAATTTGACTTGAATACTGAGA | Y2H: pB1880/81 |
| Reverse | *AthARF4* | ACCGCGGTGGCGGCCGCTCAAACCCTAGTGATTGTAGGAGA | Y2H: pB1880 |
| Reverse | *AthARF4* | ACTTACTTAGCGGCCGCTCAAACCCTAGTGATTGTAGGAGA | Y2H: pB1881 |
| Forward | *CriTPLa* | ACGACAAGGGGTCGACCATGTCGTCACTCAGTCGTGAAC | Y2H: pB1880/81 |
| Reverse | *CriTPLa* | ACCGCGGTGGCGGCCGCTCACCTTGCAGCTTGCTCAG | Y2H: pB1880 |

| | | | |
|---|---|---|---|
| Reverse | *CriTPLa* | ACTTACTTAGCGGCCGCTCACCTTGCAGCTTGCTCAG | Y2H: pB1881 |
| Forward | *CriTPLb* | ACGACAAGGGGTCGACCATGTCCTCTCTCAGCCGGGA | Y2H: pB1880/81 |
| Reverse | *CriTPLb* | ACCGCGGTGGCGGCCGCTTACTTTGGAGCCTCCAATTTCACA | Y2H: pB1880 |
| Reverse | *CriTPLb* | ACTTACTTAGCGGCCGCTTACTTTGGAGCCTCCAATTTCACA | Y2H: pB1881 |
| Forward | *CriTPLc* | ACGACAAGGGGTCGACCATGTCCTCGCTCAGCCGTG | Y2H: pB1880/81 |
| Reverse | *CriTPLc* | ACCGCGGTGGCGGCCGCCTACCTTGTGGCCTGCTCTGAG | Y2H: pB1880 |
| Reverse | *CriTPLc* | ACTTACTTAGCGGCCGCCTACCTTGTGGCCTGCTCTGAG | Y2H: pB1881 |
| Forward | *CriTPLd* | ACGACAAGGGGTCGACCATGTCATCGTTGAGTAGGGAGC | Y2H: pB1880/81 |
| Reverse | *CriTPLd* | ACCGCGGTGGCGGCCGCCTATCTTGGAGCTTGCTCTGTAAC | Y2H: pB1880 |
| Reverse | *CriTPLd* | ACTTACTTAGCGGCCGCCTATCTTGGAGCTTGCTCTGTAAC | Y2H: pB1881 |
| Forward | *CriTPLe* | ACGACAAGGGGTCGACCATGTCCTCGCTCAGCCGTG | Y2H: pB1880/81 |
| Reverse | *CriTPLe* | ACCGCGGTGGCGGCCGCCTACCTTGTGGCCTGCTCTGAG | Y2H: pB1880 |
| Reverse | *CriTPLe* | ACTTACTTAGCGGCCGCCTACCTTGTGGCCTGCTCTGAG | Y2H: pB1881 |
| Forward | *GbiTPLa* | ACGACAAGGGGTCGACCATGTCTTCCCTCAGCAGAGAGC | Y2H: pB1880/81 |
| Reverse | *GbiTPLa* | ACCGCGGTGGCGGCCGCTCACCTTGGGGTTTGCTCTG | Y2H: pB1880 |
| Reverse | *GbiTPLa* | ACTTACTTAGCGGCCGCTCACCTTGGGGTTTGCTCTG | Y2H: pB1881 |
| Forward | *GbiTPLb* | ACGACAAGGGGTCGACCATGTCTTCGCTTAGTAGAGAGCTTG | Y2H: pB1880/81 |
| Reverse | *GbiTPLb* | ACCGCGGTGGCGGCCGCTCACCTAGGACCTTGTTCTGAGC | Y2H: pB1880 |
| Reverse | *GbiTPLb* | ACTTACTTAGCGGCCGCTCACCTAGGACCTTGTTCTGAGC | Y2H: pB1881 |
| Forward | *GbiTPLc* | ACGACAAGGGGTCGACCATGTCCTCTTTAAGCAGGGAACTCG | Y2H: pB1880/81 |
| Reverse | *GbiTPLc* | ACCGCGGTGGCGGCCGCCTACCTTGGAGGTGGTTCTGAACC | Y2H: pB1880 |
| Reverse | *GbiTPLc* | ACTTACTTAGCGGCCGCCTACCTTGGAGGTGGTTCTGAACC | Y2H: pB1881 |
| Forward | *GbiTPLd* | ACGACAAGGGGTCGACCATGTCTTCTTTGAGCAGGGAACTGG | Y2H: pB1880/81 |
| Reverse | *GbiTPLd* | ACCGCGGTGGCGGCCGCCTACCTTGGAGGTAATTCTGATCCT | Y2H: pB1880 |
| Reverse | *GbiTPLd* | ACTTACTTAGCGGCCGCCTACCTTGGAGGTAATTCTGATCCT | Y2H: pB1881 |
| Forward | *GbiTPLe* | ACGACAAGGGGTCGACCATGTCTTCCCTCAGTAGGGAATTGG | Y2H: pB1880/81 |
| Reverse | *GbiTPLe* | ACCGCGGTGGCGGCCGCTTAATTGCTAGTTGGTGGTGCATG | Y2H: pB1880 |

| | | | |
|---|---|---|---|
| Reverse | *GbiTPLe* | ACTTACTTAGCGGCCGCTTAATTG CTAGTTGGTGGTGCATG | Y2H: pB1881 |
| Forward | *AtrTPR1* | ACGACAAGGGGTCGACCATGTCT TCTCTGAGCAGGGAGC | Y2H: pB1880/81 |
| Reverse | *AtrTPR1* | ACCGCGGTGGCGGCCGCCCTTG GAGGCTGCTCTGATTG | Y2H: pB1880 |
| Reverse | *AtrTPR1* | ACTTACTTAGCGGCCGCCCTTGG AGGCTGCTCTGATTG | Y2H: pB1881 |
| Forward | *AtrTPR2* | ACGACAAGGGGTCGACCATGTCT TCCTTGAGCAGGGAGC | Y2H: pB1880/81 |
| Reverse | *AtrTPR2* | ACCGCGGTGGCGGCCGCCCTCT GAGAGGGTTCAGAGGG | Y2H: pB1880 |
| Reverse | *AtrTPR2* | ACTTACTTAGCGGCCGCCCTCTG AGAGGGTTCAGAGGG | Y2H: pB1881 |
| Forward | *NthTPLa* | ACGACAAGGGGTCGACCATGTCG TCTCTTAGTAGAGAACTCG | Y2H: pB1880/81 |
| Reverse | *NthTPLa* | ACCGCGGTGGCGGCCGCTCATCT TTGCGGCTGGTC | Y2H: pB1880 |
| Reverse | *NthTPLa* | ACTTACTTAGCGGCCGCTCATCTT TGCGGCTGGTC | Y2H: pB1881 |
| Forward | *NthTPLb* | ACGACAAGGGGTCGACCATGTCG TCGCTTAGTAGAGAACTC | Y2H: pB1880/81 |
| Reverse | *NthTPLb* | ACCGCGGTGGCGGCCGCTCATCT CTGAGGCTGATCCGTG | Y2H: pB1880 |
| Reverse | *NthTPLb* | ACTTACTTAGCGGCCGCTCATCTC TGAGGCTGATCCGTG | Y2H: pB1881 |
| Forward | *NthTPR1* | ACGACAAGGGGTCGACCATGTCG TCTCTCAGTAGAGAACTCG | Y2H: pB1880/81 |
| Reverse | *NthTPR1* | ACCGCGGTGGCGGCCGCTTACCT TGGTGGCTGCTCAGAG | Y2H: pB1880 |
| Reverse | *NthTPR1* | ACTTACTTAGCGGCCGCTTACCTT GGTGGCTGCTCAGAG | Y2H: pB1881 |
| Forward | *NthTPR2a* | ACGACAAGGGGTCGACCATGTCA TCCTTAAGCAGGGAACT | Y2H: pB1880/81 |
| Reverse | *NthTPR2a* | ACCGCGGTGGCGGCCGCTCACCT GGAAGGAGGTTCTGATACC | Y2H: pB1880 |
| Reverse | *NthTPR2a* | ACTTACTTAGCGGCCGCTCACCT GGAAGGAGGTTCTGATACC | Y2H: pB1881 |
| Forward | *NthTPR2b* | ACGACAAGGGGTCGACCATGTCA TCCTTAAGCAGGGAGCTTG | Y2H: pB1880/81 |
| Reverse | *NthTPR2b* | ACCGCGGTGGCGGCCGCTCACCT GGAAGCAGGTTCTGAAG | Y2H: pB1880 |
| Reverse | *NthTPR2b* | ACTTACTTAGCGGCCGCTCACCT GGAAGCAGGTTCTGAAG | Y2H: pB1881 |
| Forward | *NthTPR3* | ACGACAAGGGGTCGACCATGTCA TCTTTAAGCAGAGAATTGG | Y2H: pB1880/81 |
| Reverse | *NthTPR3* | ACCGCGGTGGCGGCCGCTCATCT TGAAGTCTGCTCAGAGGC | Y2H: pB1880 |
| Reverse | *NthTPR3* | ACTTACTTAGCGGCCGCTCATCTT GAAGTCTGCTCAGAGGC | Y2H: pB1881 |
| Forward | *AthTPL* | ACGACAAGGGGTCGACCATGTCT TCTCTTAGTAGAGAGCTC | Y2H: pB1880/81 |
| Reverse | *AthTPL* | ACCGCGGTGGCGGCCGCTCATCT CTGAGGCTGATCAGAT | Y2H: pB1880 |

| | | | |
|---|---|---|---|
| Reverse | *AthTPL* | ACTTACTTAGCGGCCGCTCATCTCTGAGGCTGATCAGAT | Y2H: pB1881 |
| Reverse | *AthETT-GbiARF3/4* | CTGCAACTGAACCAGATGGTTCGATCTCCCATGG | Y2H Domain Swaps |
| Forward | *AthETT-GbiARF3/4* | ACCATCTGGTTCAGTTGCAGGATTGAATGTTTC | Y2H Domain Swaps |
| Reverse | *AthETT-AtrETT* | CAGACCCCAAACCAGATGGTTCGATCTCCCATGG | Y2H Domain Swaps |
| Forward | *AthETT-AtrETT* | ACCATCTGGTTTGGGGTCTGTTCCAGTATTTAGC | Y2H Domain Swaps |
| Reverse | *GbiARF3/4-AthETT* | TGGAGATGGAAATACATGGCTCAATTTCCCAT | Y2H Domain Swaps |
| Forward | *GbiARF3/4-AthETT* | GCCATGTATTTCCATCTCCAATTCAGGCAGC | Y2H Domain Swaps |
| Reverse | *GbiARF3/4-AtrETT* | CAGACCCCAAAATACATGGCTCAATTTCCCAT | Y2H Domain Swaps |
| Forward | *GbiARF3/4-AtrETT* | GCCATGTATTTTGGGGTCTGTTCCAGTATTTAGC | Y2H Domain Swaps |
| Reverse | *AtrETT-AthETT* | TGGAGATGGAAAGGTCGATCTCCCATGG | Y2H Domain Swaps |
| Forward | *AtrETT-AthETT* | GATCGACCTTTCCATCTCCAATTCAGGCAGC | Y2H Domain Swaps |
| Reverse | *AtrETT-GbiARF3/4* | CTGCAACTGAAAGGTCGATCTCCCATGG | Y2H Domain Swaps |
| Reverse | *AtrETT-GbiARF3/4* | GATCGACCTTTCAGTTGCAGGATTGAATGTTTC | Y2H Domain Swaps |
| Reverse | *AthETT-GbiARF3/4* | AAGACACAATTTCTTGACCTTGCAAGACCTTATGG | Y2H Domain Swaps |
| Reverse | *AthETT-GbiARF3/4* | AGGTCAAGAAATTGTGTCTTTGAAGGCACCCC | Y2H Domain Swaps |
| Reverse | *AthETT-AtrETT* | AATGGAAAATTTCTTGACCTTGCAAGACCTTATGG | Y2H Domain Swaps |
| Forward | *AthETT-AtrETT* | AGGTCAAGAAATTTTCCATTTGAAATCACAGAACA | Y2H Domain Swaps |
| Forward | *pAthETT* | GAGCTCGGTACCCGGGGATCCGAGCAATCCTATACGGAGTTCT | Plasmid –Promoter |
| Reverse | *pAthETT-MpoARF2* | CTTCTGACATTAAAGAGAGAGAAACAGAGATAAAG | Promoter –CDS |
| Reverse | *pAthETT-CriARF3/4* | CGAGATGCATTAAAGAGAGAGAAACAGAGATAAAG | Promoter –CDS |
| Reverse | *pAthETT-GbiARF3/4* | CAATTTCCATTAAAGAGAGAGAAACAGAGATAAAG | Promoter –CDS |
| Reverse | *pAthETT-AtrETT* | CAATGCCCATTAAAGAGAGAGAAACAGAGATAAAG | Promoter –CDS |
| Reverse | *pAthETT-AtrARF4* | CAATTTCCATTAAAGAGAGAGAAACAGAGATAAAG | Promoter –CDS |
| Reverse | *pAthETT-NthETT* | CGATCTCCATTAAAGAGAGAGAAACAGAGATAAAG | Promoter –CDS |
| Reverse | *pAthETT-NthARF4* | CAATTTCCATTAAAGAGAGAGAAACAGAGATAAAG | Promoter –CDS |
| Reverse | *pAthETT-AthETT* | AACCACCCATTAAAGAGAGAGAAACAGAGATAAAG | Promoter –CDS |
| Reverse | *pAthETT-AthARF4* | CAAATTCCATTAAAGAGAGAGAAACAGAGATAAAG | Promoter –CDS |

| | | | |
|---|---|---|---|
| Forward | *pAthETT-MpoARF2* | TCTCTCTTTAATGTCAGAAGCATCTTCCATCAC | Promoter –CDS |
| Forward | *pAthETT-CriARF3/4* | TCTCTCTTTAATGCATCTCGACTCGCAGC | Promoter –CDS |
| Forward | *pAthETT-GbiARF3/4* | TCTCTCTTTAATGGAAATTGATCTCAACAGTCC | Promoter –CDS |
| Forward | *pAthETT-AtrETT* | TCTCTCTTTAATGGGCATTGATCTGAACCGG | Promoter –CDS |
| Forward | *pAthETT-AtrARF4* | TCTCTCTTTAATGGAAATTGATCTCAACTGCG | Promoter –CDS |
| Forward | *pAthETT-NthETT* | TCTCTCTTTAATGGAGATCGATCTAAACAGGGTCG | Promoter –CDS |
| Forward | *pAthETT-NthARF4* | TCTCTCTTTAATGGAAATTGATCTCAACAGTGC | Promoter –CDS |
| Forward | *pAthETT-AfiETT* | TCTCTCTTTAATGGAAATCGATCTGAACGCCA | Promoter –CDS |
| Forward | *pAthETT-AfiARF4* | CAATTTCCATTAAAGAGAGAGAAACAGAGATAAAG | Promoter –CDS |
| Forward | *pAthETT-AthETT* | TCTCTCTTTAATGGGTGGTTTAATCGATCTGAACG | Promoter –CDS |
| Forward | *pAthETT-AthARF4* | TCTCTCTTTAATGGAATTTGACTTGAATACTGAGA | Promoter –CDS |
| Reverse | *MpoARF2* | ATGGTCTTTGTAGTCAAGCTTCTACATGTCGTCGCC | Plasmid – CDS |
| Reverse | *CriARF3/4* | ATGGTCTTTGTAGTCAAGCTTTTAACCTGAGATGGA | Plasmid – CDS |
| Reverse | *GbiARF3/4* | ATGGTCTTTGTAGTCAAGCTTTTAAATTCTTGTTGCTGGTGGAG | Plasmid – CDS |
| Reverse | *AtrETT* | ATGGTCTTTGTAGTCAAGCTTTTACACAGCTCTAGCAAGGCC | Plasmid – CDS |
| Reverse | *AtrARF4* | ATGGTCTTTGTAGTCAAGCTTCTATTCGAGTCCTCTTGTTAT | Plasmid – CDS |
| Reverse | *NthETT* | ATGGTCTTTGTAGTCAAGCTTTCAAGTCCTTGTTAC | Plasmid – CDS |
| Reverse | *NthARF4* | ATGGTCTTTGTAGTCAAGCTTCTAGATGTTGTTTTC | Plasmid – CDS |
| Reverse | *AthETT* | ATGGTCTTTGTAGTCAAGCTTCTAGAGAGCAATGTC | Plasmid – CDS |
| Reverse | *AthARF4* | ATGGTCTTTGTAGTCAAGCTTTCAAACCCTAGTGATTGTA | Plasmid – CDS |
| Reverse | *AthETT-GbiARF3/4* | ATGGTCTTTGTAGTCAAGCTTTTAAGGCCTCGTTACTGTCAAAG | Domain Swaps |
| Reverse | *AthETT-GbiARF3/4* | ATGGTCTTTGTAGTCAAGCTTACGCCCCAAAGCTGGAACT | Domain Swaps |
| Reverse | *AthETT-AtrETT* | ATGGTCTTTGTAGTCAAGCTTTCAAACCCTAGTGATTGTA | Domain Swaps |
| Reverse | *AthETT-AtrETT* | ATGGTCTTTGTAGTCAAGCTTGCGGTTGGCTGTTTTATTCA | Domain Swaps |
| Reverse | *AthETT* | ATGGTCTTTGTAGTCAAGCTTGAGAGCAATGTCTAGCAACATGTC | Domain Swaps |
| Forward | *arf4$^{GE}$* | AACAAGCACCTCCATTTAG | Genotyping – WT/mutant |
| Reverse | *arf4$^{GE}$* | TCCATTTACTCTGAGCTTTG | Genotyping – WT |
| Reverse | *arf4$^{GE}$* | ACACATCCCATAAACTCACA | Genotyping –mutant |
| Forward | *pAthETT* | AGAGGGTAAAAGTCATGAAG | Genotyping |

| | | | |
|---|---|---|---|
| | *5'UTR* | | |
| Reverse | *pCambia1305* | TCACTTATCGTCATCGTCCT | Genotyping |
| Reverse | *MpoARF2* | CTTTGGGTTGCAGTTGTT | Genotyping |
| Reverse | *CriARF3/4* | ACGATACTTTTGATGGGGA | Genotyping |
| Forward | *GbiARF3/4* | GTGATTACCAGCAGCCTTTT | Genotyping |
| Forward | *AtrETT* | CGTTTTCAGACTTTGGGGAA | Genotyping |
| Forward | *AtrARF4* | TTTTCTACAACCCAAGGGCA | Genotyping |
| Forward | *NthETT* | TGAATGCCAGAGTGTAGA | Genotyping |
| Reverse | *NthARF4* | CATCCCAACGTACCATCA | Genotyping |
| Reverse | *AthETT* | CCCTCGCTTTAAATCTCATC | Genotyping |
| Reverse | *AthARF4* | AACAAGCACCTCCATTTAG | Genotyping |
